# Immune dysregulation in SHARPIN-deficient mice is dependent on CYLD-mediated cell death

**DOI:** 10.1101/2020.01.27.919076

**Authors:** Rosalind L. Ang, John P. Sundberg, Shao-Cong Sun, Virginia L. Gillespie, Peter S. Heeger, Huabao Xiong, Sergio A. Lira, Adrian T. Ting

**Affiliations:** Precision Immunology Institute, Icahn School of Medicine at Mount Sinai, New York, NY 10029, USA; The Jackson Laboratory, Bar Harbor, ME 04609; Department of Immunology, MD Anderson Cancer Center, The University of Texas, Houston, TX 77030; Center for Comparative Medicine and Surgery, Icahn School of Medicine at Mount Sinai, New York, NY 10029, USA; Department of Medicine, Translational Transplant Research Center, Recanati Miller Transplant Institute, Icahn School of Medicine at Mount Sinai, New York, NY 10029, USA; Tisch Cancer Institute, Department of Medicine, Icahn School of Medicine at Mount Sinai, New York, NY 10029, USA; Department of Immunology, Mayo Clinic, Rochester, MN 55905, USA

## Abstract

SHARPIN, together with RNF31/HOIP and RBCK1/HOIL1, form the linear ubiquitin chain assembly complex (LUBAC) E3 ligase that catalyzes M1-linked poly-ubiquitination. Mutations in *RNF31/HOIP* and *RBCK/HOIL1* in humans and *Sharpin* in mice lead to auto-inflammation and immunodeficiency but the mechanism underlying the immune dysregulation remains unclear. We now show that the phenotype of the *Sharpin^-/-^* mice is dependent on CYLD, the deubiquitinase that removes K63-linked poly-ubiquitin chains. The dermatitis, disrupted splenic architecture, and loss of Peyer’s patches in the *Sharpin^-/-^* mice were fully reversed in *Sharpin^-/-^Cyld^-/-^* mice. There is enhanced association of RIPK1 with the death-inducing signaling complex (DISC) following TNF stimulation in *Sharpin^-/-^* cells, and this is dependent on CYLD since it is reversed in *Sharpin^-/-^Cyld^-/-^* cells. Enhanced RIPK1 recruitment to the DISC in *Sharpin^-/-^* cells correlated with impaired phosphorylation of CYLD at serine 418, a modification reported to inhibit its enzymatic activity. The dermatitis in the *Sharpin^-/-^* mice was also ameliorated by the conditional deletion of *Cyld* using *LysM-cre* or *Cx3cr1-cre* indicating that CYLD-dependent death of myeloid cells is inflammatory. Our studies reveal that under physiological conditions, TNF- and RIPK1-dependent cell death is suppressed by the linear ubiquitin-dependent inhibition of CYLD. The *Sharpin^-/-^* phenotype illustrates the pathological consequences when CYLD inhibition fails.

**Short Summary:** In the absence of SHARPIN, cells fail to properly regulate the deubiquitinase CYLD, leading to RIPK1-mediated cell death. Deletion of *Cyld* reverses the sensitivity of *Sharpin^-/-^* cells to TNF-induced cell death, as well as the multi-organ inflammation and immune dysfunction observed in *Sharpin^-/-^* mice.

## Introduction

Linear or M1-linked poly-ubiquitin modifications play a major role in regulating immune responses (1–3). This modification is catalyzed by the linear ubiquitin chain assembly complex (LUBAC) E3 ligase, which consists of the catalytic component RNF31/HOIP in complex with two other essential proteins with non-catalytic functions RBCK1/HOIL1 and SHARPIN (4–10). Linear ubiquitination plays a key role in signaling downstream of several immune receptors including Tumor Necrosis Factor Receptor Superfamily (TNFRSF) members, antigen receptors and pattern recognition receptors (1, 11). In the case of TNFα (TNF), LUBAC components are rapidly recruited to TNFR1 following receptor ligation to ubiquitinate target molecules, providing scaffolding for the formation of multi-molecular complexes that propagate signaling. One critical signaling complex assembled by linear ubiquitin is the I-κB kinase (IKK) complex that phosphorylates I-κBα (5, 6, 12), a step required for subsequent degradation of I-κBα and induction of NF-κB-dependent gene transcription. NEMO (encoded by the *Ikbkg* gene), a structural component of the IKK complex, is modified by linear ubiquitination and also possesses a linear ubiquitin-binding domain that facilitates oligomerization of the IKK complex (5, 6, 12, 13). Cells deficient in linear ubiquitination have impaired IKK activity and NF-κB signaling in response to TNF stimulation (5, 7–9). Since IKK can phosphorylate substrates other than I-κB (14–16), whether linear ubiquitin regulates NF-κB-independent pathways and their downstream biological processes remains poorly understood.

Consistent with its role in the signaling of immune receptors, LUBAC components are critical for proper immune regulation. Loss-of-function mutations in *RNF31/HOIP* and in *RBCK1/HOIL1* in humans cause immunodeficiency, autoinflammation, amylopectinosis, and lymphangiectasia (17, 18). In mice, a loss-of-function mutation in *Sharpin* has been found in the chronic proliferative dermatitis (*cpdm*) mice (hereafter referred to as *Sharpin^-/-^*), which spontaneously develops dermatitis, multi-organ inflammation and displays immunodeficiency (7-9, 19-21). Crossing of *Sharpin^-/-^* mice with *Tnf^-/-^* mice reversed the skin inflammation indicating that some of the lesion in this mutant mouse are TNF-driven (9). *Sharpin^-/-^* cells exhibited enhanced cell death in response to TNF (8, 9) suggesting that the phenotype of the *Sharpin^-/-^* mice could be due to inappropriate cell death. Subsequent genetic crosses with knockouts of death-signaling molecules provide further support for this hypothesis. Deletion of the necroptosis-signaling molecule *Ripk3* delayed but did not prevent the dermatitis but additional deletion of a single allele of the apoptosis-signaling molecule *Casp8* or skin-specific deletion of *Fadd* reversed the dermatitis in the *Sharpin^-/-^* mice (22, 23). Furthermore, crossing with a kinase-inactive *Ripk1^tm1.1Gsk^* (*Ripk1^K45A^*) allele also reversed the phenotype of the *Sharpin^-/-^* mice (24). These genetic analyses suggest that the phenotype of the *Sharpin^-/-^* mouse is due to excessive RIPK1-dependent cell death and we recently suggested the term *ripoptocide* to denote this manner of cell death orchestrated by RIPK1 (25). One mechanism by which *Sharpin* deficiency could lead to aberrant RIPK1 death-signaling is impaired phosphorylation of RIPK1 catalyzed by members of the IKK family, which normally inhibits its death-promoting activity (26–30). Whether the *Sharpin* deficiency also affects other molecules involved in regulating cell death is unclear.

Another form of poly-ubiquitin modification, one in which the covalent linkage occurs at K63 of the ubiquitin molecule, is also critical in TNF signaling (31, 32). This form of ubiquitin modification is mediated by TRAF2-dependent recruitment of the cIAP1/2 E3 ligases (encoded by *Birc2* and *Birc3*) and one key molecule modified in this manner is RIPK1. Cells in which the ubiquitin acceptor site of RIPK1 at K377 was mutated were more sensitive to TNF-mediated apoptosis, similar to cells in which TRAF2 and cIAP1/2 were inhibited (33–35). Cells treated with Second Mitochondria-Derived Activator Of Caspase (SMAC) mimetics, which induce the degradation and loss of cIAP1/2, displayed reduced poly-ubiquitination of RIPK1 (34, 35). This cell protective function of K63-linked poly-ubiquitination of RIPK1 is initially transcriptional-independent (i.e., does not require NF-κB) but subsequently feeds into NF-κB-dependent induction of pro-survival genes. A unifying model proposes that there are two cell death checkpoints in the TNFR1 signaling pathway. The early checkpoint (Checkpoint 1) prevents RIPK1 from becoming a death-signaling molecule via K63-linked poly-ubiquitination, which is followed by the NF-κB-dependent induction of pro-survival genes during the late checkpoint (Checkpoint 2) (36–38). Disruption of Checkpoint 1 is often achieved *in vitro* by blocking RIPK1 ubiquitination using SMAC mimetics, which leads to CASP8-dependent apoptosis or RIPK3/MLKL-dependent necroptosis (35, 39–41). The relationship between linear and K63 ubiquitin linkages in regulating cell death and potentially in the immune dysfunction observed in LUBAC deficiencies is currently unclear.

We now report that the molecule CYLD is central to the immunopathology observed in the *Sharpin^-/-^* mice. CYLD is a deubiquitinase that dismantles K63-linked poly-ubiquitin chains (42) and its activity was shown to be suppressed by IKKβ-mediated phosphorylation (43). CYLD has also been reported to associate with RNF31/HOIP via SPATA2 (44–48). We found that SHARPIN-deficient cells, which are deficient in IKK activity, display reduced CYLD phosphorylation following receptor ligation. Coincident with defective CYLD phosphorylation in the SHARPIN-deficient cells, there is enhanced association of its substrate RIPK1 with a death-inducing signaling complex (DISC) to induce apoptosis and necroptosis, which is reversed in SHARPIN and CYLD double-deficient cells. The phenotype of the *Sharpin^-/-^* mice is reversed by a compound deletion of *Cyld* and importantly, conditional deletion of *Cyld* in myeloid cells significantly reversed the dermatitis. Our study provides evidence that linear ubiquitin-dependent suppression of CYLD is one mechanism that keeps ripoptocide at bay and to maintain immune homeostasis, disruption of which leads to immune dysfunction.

## Results

### *Cyld* is essential for the development of inflammation in *Sharpin*-deficient mice

*Sharpin^-/-^* cells exhibited heightened sensitivity to TNF-induced killing (9). Since CYLD has been reported to be required for TNF to induce RIPK1-dependent apoptosis and necroptosis (34, 49–53), we asked if CYLD has any role in the sensitivity of *Sharpin^-/-^* cells to TNF-induced cell death. We first confirmed that *Sharpin^-/-^* mouse embryonic fibroblasts (MEF) were more sensitive to TNF-induced necroptosis than their wild type counterparts, an effect that was reversed when the *Sharpin^-/-^* MEF were complemented with *Sharpin* but not with a control gene (Supplementary Figure S1A). Comparison of the *Sharpin^-/-^* MEF complemented with *Sharpin* versus the control gene showed SHARPIN-deficient cells to be more sensitive to TNF-induced necroptosis in a dose and time-dependent manner (Supplementary Figure S1B-D). Furthermore, when both SHARPIN-sufficient and SHARPIN-deficient MEF were stably transfected with the non-degradable IκB super-repressor (IκBSR) to block NF-κB induction, SHARPIN-deficient MEF remained more sensitive to death compared to SHARPIN-sufficient MEF (Supplementary Figure S1E). Blotting of nuclear extracts from the *Sharpin*-complemented MEFs transfected with the IκBSR confirmed the effectiveness of the IκBSR in blocking translocation of the NFκB p65 subunit to the nucleus (Supplementary Figure S1F). This indicated that the loss of SHARPIN could sensitize cells to death through a NF-κB-independent mechanism, reminiscent of the effect of mutating the acceptor site on RIPK1 for K63-linked poly-ubiquitin (33). As an initial test of a potential role for CYLD, we examined the effect of knocking down CYLD on necroptosis in SHARPIN-deficient MEF. Knocking down CYLD significantly reduced the level of necroptosis in response to TNF (Supplementary Figure S1G & H). This observation suggested that the loss of SHARPIN led to CYLD-dependent death.

These results prompted us to test whether CYLD regulates the phenotype of the *Sharpin^-/-^* mice. To test this hypothesis, we generated *Sharpin^-/-^Cyld^-/-^* double deficient animals. Strikingly, the loss of *Cyld* fully prevented the spontaneous development of dermatitis that develops in *Sharpin^-/-^* mice (Figure 1A &B). Histological analysis showed that ulceration, thickening of the epidermis and leukocytic infiltration observed in *Sharpin^-/-^* skin were absent in *Sharpin^-/-^Cyld^-/-^* skin (DKO, Figure 1C). Whereas *Sharpin^-/-^* mice were smaller and weighed less than age-matched wild type mice, sizes and weights of the *Sharpin^-/-^Cyld^-/-^* mice and the wild type controls did not differ (Figure 1A, Supplementary Figure S2A). The skin of *Sharpin^-/-^* mice were reported to be infiltrated by immune cells including granulocytes and macrophages (19). Immunohistochemical analysis confirmed the presence of CD45^+^ hematopoietic cells in the skin from *Sharpin^-/-^* mice but not from *Sharpin^-/-^Cyld^-/-^* mice (Figure 1D & E). F4/80+ macrophages were also detected in the inflamed skin of the *Sharpin^-/-^* mice (Figure 1F & G). Thus, the dermatitis in the *Sharpin^-/-^* mice was ameliorated by the compound loss of *Cyld*.

**Figure 1.**
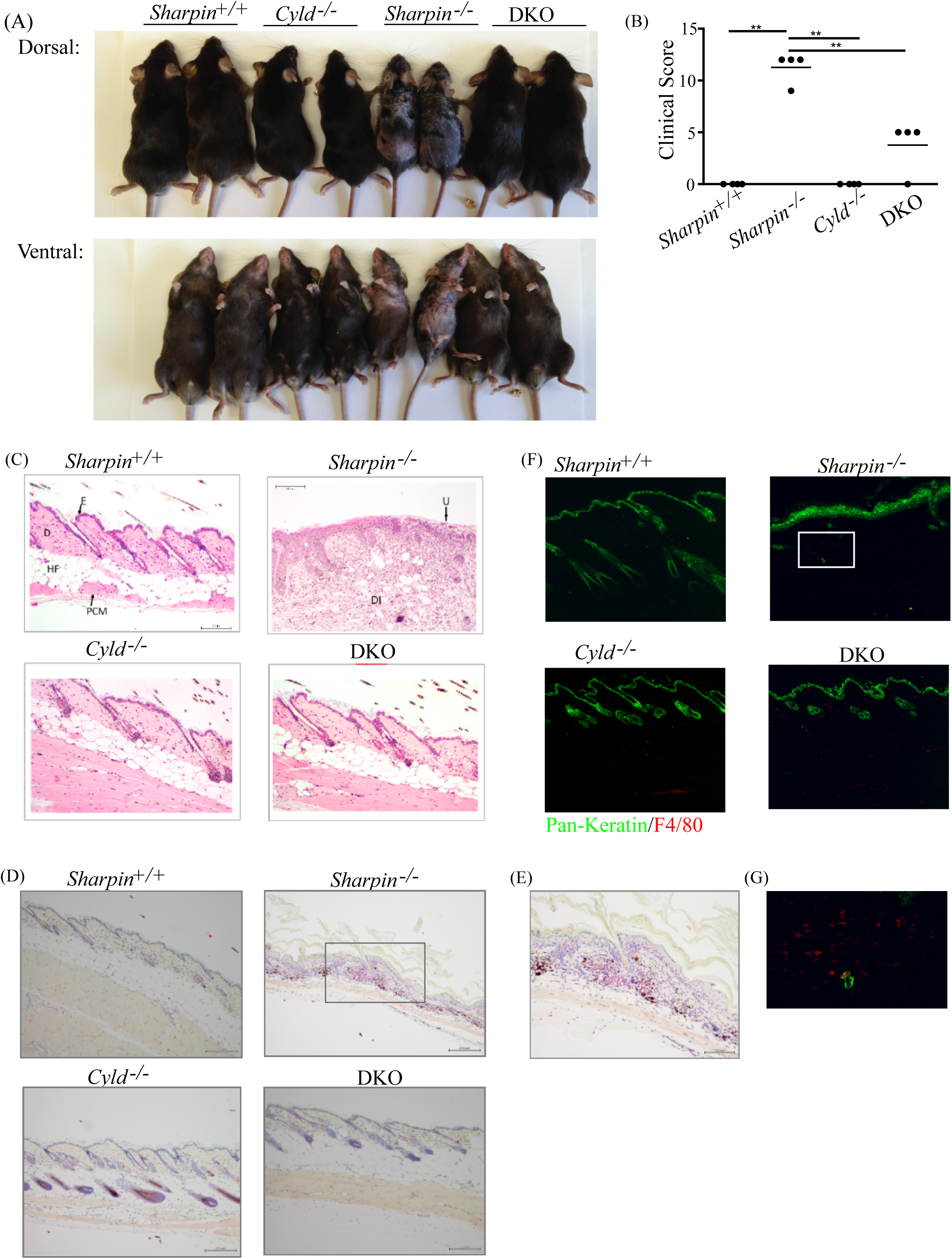
*Cyld* is essential for the development of inflammation in *Sharpin^-/-^* mice. (A) Representative 16-week-old female *Sharpin^+/+^*, *Cyld^-/-^*, *Sharpin^-/-^*, and *Sharpin^-/-^ Cyld^-/-^* (DKO) mice. (B) A cohort of 16-week-old female mice was examined by a pathologist in a double-blind fashion. The clinical score of mice from the indicated genotypes are shown (n=4 per genotype). One-way ANOVA analysis was performed. *P<0.05, **P<0.01. (C) Histology of skin taken from mice in (B) by H&E staining (100X). Representative photomicrographs from each genotype are shown. E: epidermis; D: dermis; HF: hypodermal fat; PCM: panniculus carnosus muscle; U: ulcerated area with complete loss of epidermis, replacement by necrosis and inflammation; DI: dermal inflammation. (D) Skin sections taken from mice in (B) were analyzed by immunohistochemistry with anti-CD45 to label hematopoietic cells (100X). CD45+ cells are stained red. Representative photomicrographs from each genotype are shown. (E) Rectangular insert of the *Sharpin^-/-^* skin section from panel (D) above is enlarged for visualization of the CD45+ cells. (F) Skin sections taken from mice in (B) were analyzed by immunofluorescence with anti-F4/80 to detect macrophages (100X). Representative photomicrographs from each genotype are shown. (g) Rectangular insert of the *Sharpin^-/-^* skin section from panel (F) above is enlarged for visualization of F4/80+ macrophages.

Lymphoid organs in *Sharpin^-/-^* mice are known to be abnormal (20, 21). Gross examination of immune organs showed that the enlarged spleen, shrunken thymus and mesenteric lymph nodes and the enlarged liver typical of *Sharpin^-/-^* mice were absent in the *Sharpin^-/-^Cyld^-/-^* mice (Figure 2A, Supplementary Figure S2B & C). We observed no alterations in the weight of the hearts or the length of the colon in any of the genotypes (Supplementary Figure S2D & E). The splenic architecture of the *Sharpin^-/-^* spleen lacks lymphoid follicles whereas the *Sharpin^-/-^Cyld^-/-^* spleen is similar to that of *Sharpin^+/+^* spleen and displays no overt defect (Figure 2B). Adult *Sharpin^-/-^* mice have no Peyer’s patches in their small intestine (20, 54) but these were present in the *Sharpin^-/-^Cyld^-/-^* mice (Figure 2C & D). Steady state serum immunoglobulin levels are also perturbed in the *Sharpin^-/-^* mice (20). They have lower IgG and IgA and these abnormalities were reversed in the *Sharpin^-/-^Cyld^-/-^* mice (Figure 2E). Serum IgE level was elevated in the *Sharpin^-/-^* mice but was reversed in the *Sharpin^-/-^Cyld^-/-^* mice (Figure 2E). Consistent with the lack of Peyer’s patches, adult *Sharpin^-/-^* mice have no fecal IgA (20), but this was also reversed in the *Sharpin^-/-^Cyld^-/-^* mice (Figure 2F). Whereas several cytokines, including IFNG, IL5 and IL12/23p40 were elevated in the serum of *Sharpin^-/-^* mice compared to wild type controls, these elevations were abrogated in the *Sharpin^-/-^Cyld^-/-^* mice (Figure 2F). More modest changes in serum IL2 and IL6, but not eotaxin and CCL3 (MIP-1α), were observed in the *Sharpin^-/-^* mice (Supplementary Figure S2F-I). Together, these data demonstrate that CYLD is critical for eliciting the multi-organ inflammation and abnormal lymphoid tissues observed in the *Sharpin^-/-^* mice, suggesting dysregulated CYLD activity in the absence of SHARPIN.

**Figure 2.**
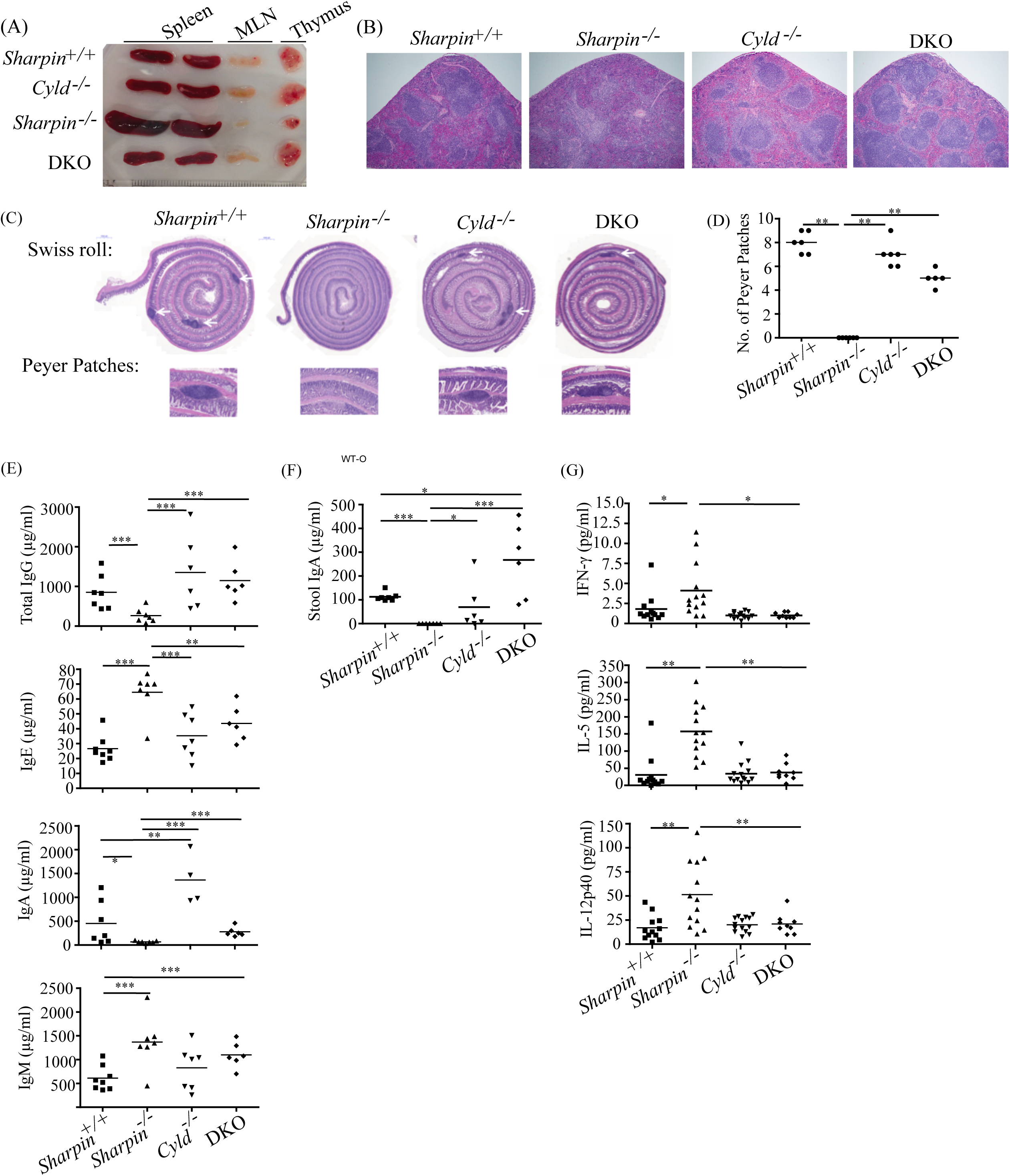
*Cyld* deletion reverses lymphoid tissue defects in *Sharpin^-/-^* mice. (A) Gross analysis of spleens, mesenteric lymph nodes (MLN) and thymi obtained from 16-week-old mice. Spleens from two individual mice of each genotype are shown. (B) Histology of spleens taken from mice of the indicated genotypes by H&E staining (100X). Representative photomicrographs from each genotype are shown. (C) H&E staining of Swiss rolls of small intestines obtained from 16 week-old mice of the indicated genotypes. White arrows indicate Peyer’s patches. Images of individual Peyer’s patches are shown below. Note *Sharpin*^-/-^ mice lack Peyer’s patches. (D) The number of Peyer’s patches was enumerated in each mouse of the different genotypes. One-way ANOVA analysis was performed. *P<0.05, **P<0.01. (E) Level of serum IgG, IgA, IgE and IgM from 10-16-week-old mice were measured by Luminex Multiplex assays. Each data point is from an individual mouse (n=6-8 mice per genotype). Error bars represent mean±SD and Student’s t-test was performed. *P<0.05, **P<0.01, ***P<0.001. (F) Stool IgA level in mice from the indicated genotypes were analyzed by ELISA. Each data point is from an individual mouse (n=6-8 mice per genotype). Error bars represent mean±SD and Student’s t-test was performed. *P<0.05, **P<0.01, ***P<0.001. (G) Level of serum IFNG, IL5 and IL12/23p40 from 10-16-week-old mice were measured by Luminex Multiplex assays. Each data point is from an individual mouse (n=9-12 mice per genotype). Error bars represent mean±SD and Student’s t-test was performed. *P<0.05, **P<0.01.

### CYLD phosphorylation is disrupted by SHARPIN deficiency

The above results showed a clear genetic interaction between SHARPIN and CYLD but the mechanistic relationship between the two molecules was unknown. Following TNFR1 ligation, an array of signaling scaffold and effector molecules such as LUBAC, cIAP1/2, TRAF2, and IKK are recruited to TNFR1. This receptor complex, also known as Complex I (55), propagates a survival signal. CYLD can also be recruited to the receptor complex via a SPATA2-dependent association with RNF31/HOIP (44–48). But how defects in linear ubiquitin could lead to CYLD activation was not known. CYLD is regulated by post-translational mechanisms, and cleavage by CASP8 and MALT1 have been reported to regulate its function (53, 56). Another potential regulatory mechanism may be phosphorylation. CYLD phosphorylation at a cluster of serine residues around Ser 418 carried out by IKKβ in a stimulus dependent fashion is known to suppress its enzymatic activity (43). Phosphorylation of CYLD at Ser 418 by IKKεwas also reported to have the same effect (57). Since SHARPIN-deficient cells were reported to have diminished IKK activity (8), we postulated that SHARPIN deficiency would also lead to diminished CYLD phosphorylation, which would result in a more active CYLD to initiate cell death. To test this, we examined the phosphorylation kinetics of CYLD in primary adult dermal fibroblast (ADF). Using an antibody that detects phosphorylated Ser 418 of CYLD, we observed an induction in CYLD phosphorylation in *Sharpin^+/+^* cells but this phosphorylation was diminished in *Sharpin^-/-^* cells (Figure 3A). To confirm that the defect in CYLD regulation is caused specifically by SHARPIN deficiency, we examined the SHARPIN-complemented MEF described in Supplementary Figure S1. Complementation of *Sharpin^-/-^* cells with *Sharpin*, but not with a control gene, restored CYLD phosphorylation (Figure 3B). Consistent with the previous study that IKKβ phosphorylates CYLD (43), blockade of IKKβ with the highly selective chemical inhibitor [5-(*p*-Fluorophenyl)-2-ureido]thiophene-3-carboxamide (TPCA-1) resulted in reduced CYLD phosphorylation in *Sharpin^+/-^* MEF, similar to that observed in *Sharpin^-/-^* MEF (Figure 3C). To extend our observations beyond fibroblasts, we examined splenic B cells. *Sharpin^-/-^* B cells also showed diminished CYLD phosphorylation when compared to their wild type counterparts (Figure 3D) though there appears to be some reduction in the level of CYLD in the *Sharpin^-/-^* B cells. Similarly, in total splenocytes, CYLD phosphorylation was detected in *Sharpin^+/+^* but not in *Sharpin^-/-^* cells (Figure 3E). Interestingly, blotting for CYLD revealed a significant loss in full-length CYLD in the *Sharpin^-/-^* splenocytes, accompanied by the detection of smaller fragments of CYLD. CASP8 and MALT1 have been shown to cleave CYLD (53, 56, 58). A 25 kD and 35 kD N-terminal fragment of CYLD that are products of CASP8- and MALT1-mediated proteolysis, respectively, are detected in the *Sharpin^-/-^* splenocytes. These observations suggest that compensatory mechanisms to remove CYLD may be invoked in the face of SHARPIN deficiency to attain a cellular state akin to deleting *Cyld* in the *Sharpin^-/-^Cyld^-/-^* mice. These findings indicate that in *Sharpin* deficiency, CYLD phosphorylation is impaired resulting in its initiation of cell death and that some cell types may attempt to avoid this fate by removing CYLD through other mechanisms.

**Figure 3.**
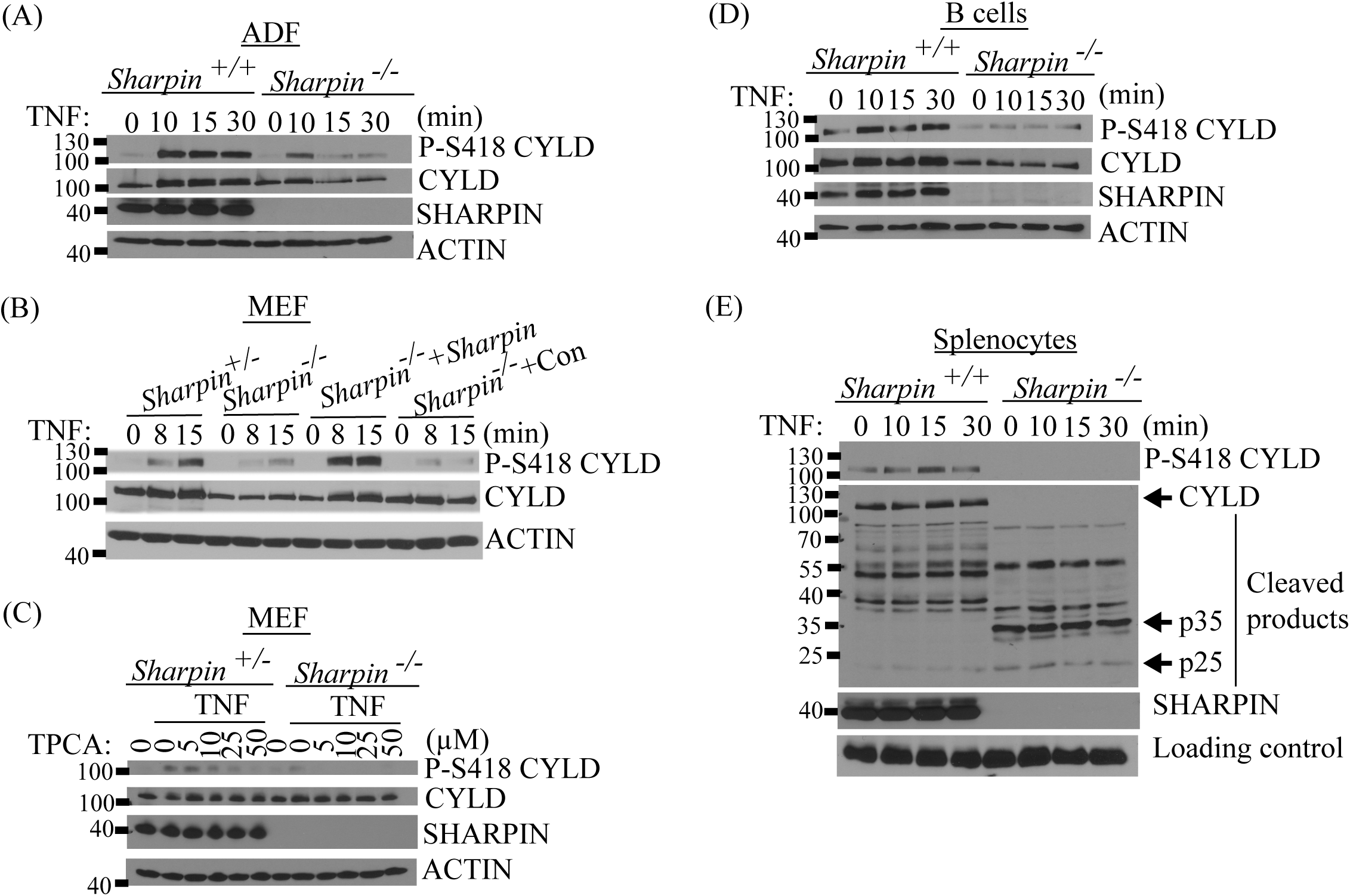
CYLD phosphorylation is impaired by SHARPIN deficiency. (A) Adult dermal fibroblasts (ADF) from *Sharpin^+/+^* and *Sharpin^-/-^* mice were treated with mTNF (100 ng/ml) for the indicated minutes. CYLD phosphorylation was detected by blotting with anti-phospho-CYLD (Ser 418), followed by blotting for total CYLD, SHARPIN and β-ACTIN. (B) *Sharpin^+/-^*, *Sharpin^-/-^* and *Sharpin^-/-^* MEF complemented with *Sharpin* or control gene were stimulated with mTNF (100 ng/ml) for the indicated times. CYLD phosphorylation was analyzed as in (A). (C) *Sharpin^+/-^*, and *Sharpin^-/-^* MEF were stimulated with 100 ng/ml mTNF for 10 min in the presence of 0 to 50 μM TPCA-1, a highly selective inhibitor against IKKβ. CYLD phosphorylation was analyzed as in (A). (D) Purified B cells from *Sharpin^+/+^* and *Sharpin^-/-^* mice were treated with mTNF (100 ng/ml) for the indicated minutes. CYLD phosphorylation was analyzed as in (A). (E) Total splenocytes from *Sharpin^+/+^* and *Sharpin^-/-^* mice were treated with mTNF (100 ng/ml) for the indicated minutes. CYLD phosphorylation was analyzed as in (A). The indicated 25 kD and 35 kD fragments of CYLD correspond to N-terminal fragments from proteolysis by CASP8 and MALT1, respectively. Data in panels A-E are representative of at least three independent experiments conducted for each panel.

### SHARPIN-deficient cells are more susceptible to CYLD-mediated cell death

Since our earlier result in MEF showed a role for SHARPIN and CYLD in regulating cell death (Supplementary Figure S1), we examined the sensitivity of ADF obtained from *Sharpin^+/+^*, *Sharpin^-/-^*, *Cyld^-/-^* and *Sharpin^-/-^Cyld^-/-^* mice to TNF-induced cell death. *Sharpin^-/-^* ADF were more susceptible to TNF-induced apoptosis and necroptosis when compared to their *Sharpin^+/+^* counterpart (Figure 4A). The increased susceptibility to both forms of death was abolished in *Sharpin^-/-^Cyld^-/-^* cells (Figure 4A), consistent with observations obtained earlier by knocking down CYLD (Supplementary Figure S1). Similarly, *Sharpin^-/-^* ADF exhibited enhanced apoptotic markers (i.e., cleaved CASP8, CASP3 and PARP) and a necroptotic marker (i.e., phospho-MLKL) when compared to *Sharpin^+/+^* ADF, and these death signatures were diminished in the *Sharpin^-/-^Cyld^-/-^* ADF (Figure 4B-E). Similar data were also observed in MEF. *Sharpin^-/-^* MEF were more sensitive to apoptosis and necroptosis, which was reversed in the *Sharpin^-/-^Cyld^-/-^* MEF (Supplementary Figure S3A-E). These results demonstrate that enhanced sensitivity to cell death in SHARPIN-deficient cells is dependent on CYLD. CYLD initiates death-signaling by removing K63-linked ubiquitin chains from RIPK1 thereby converting RIPK1 from a survival-signaling to a death-signaling effector (31, 53, 59). This conversion is biochemically detected by the translocation of RIPK1 to the FADD/CASP8 death-inducing signaling complex (DISC). We therefore sought to confirm that there was more conversion of RIPK1 to its death-signaling form in SHARPIN-deficient cells and that this was dependent on CYLD. In ADF treated with TNF, in the presence of a combination of cycloheximide and zVAD-fmk to stabilize the DISC, RIPK1 was detected in the FADD pulldowns of *Sharpin^-/-^* ADF more rapidly than in *Sharpin^+/+^* cells (Figure 4F). RIPK1 translocation to the DISC was abrogated in *Sharpin^-/-^Cyld^-/-^* ADF (Figure 4F) demonstrating that CYLD is necessary for the conversion of RIPK1 to a death-signaling molecule when SHARPIN is absent. The dependence on CYLD for RIPK1 translocation to the DISC was also observed in MEF (Supplementary Figure S3F). We confirmed expression, or lack therefore, of SHARPIN and CYLD in the MEF of the indicated genotype (Supplementary Figure S3G). As an additional confirmation, we reconstituted *Sharpin^-/-^Cyld^-/-^* MEF with retroviral vectors encoding SHARPIN only, CYLD only or both molecules together (Supplementary Figure S3H & 3I). These reconstituted MEF showed similar data (Supplementary Figure S3H) to that observed previously in ADF (Figure 4F) and MEF (Supplementary Figure S3F).

**Figure 4.**
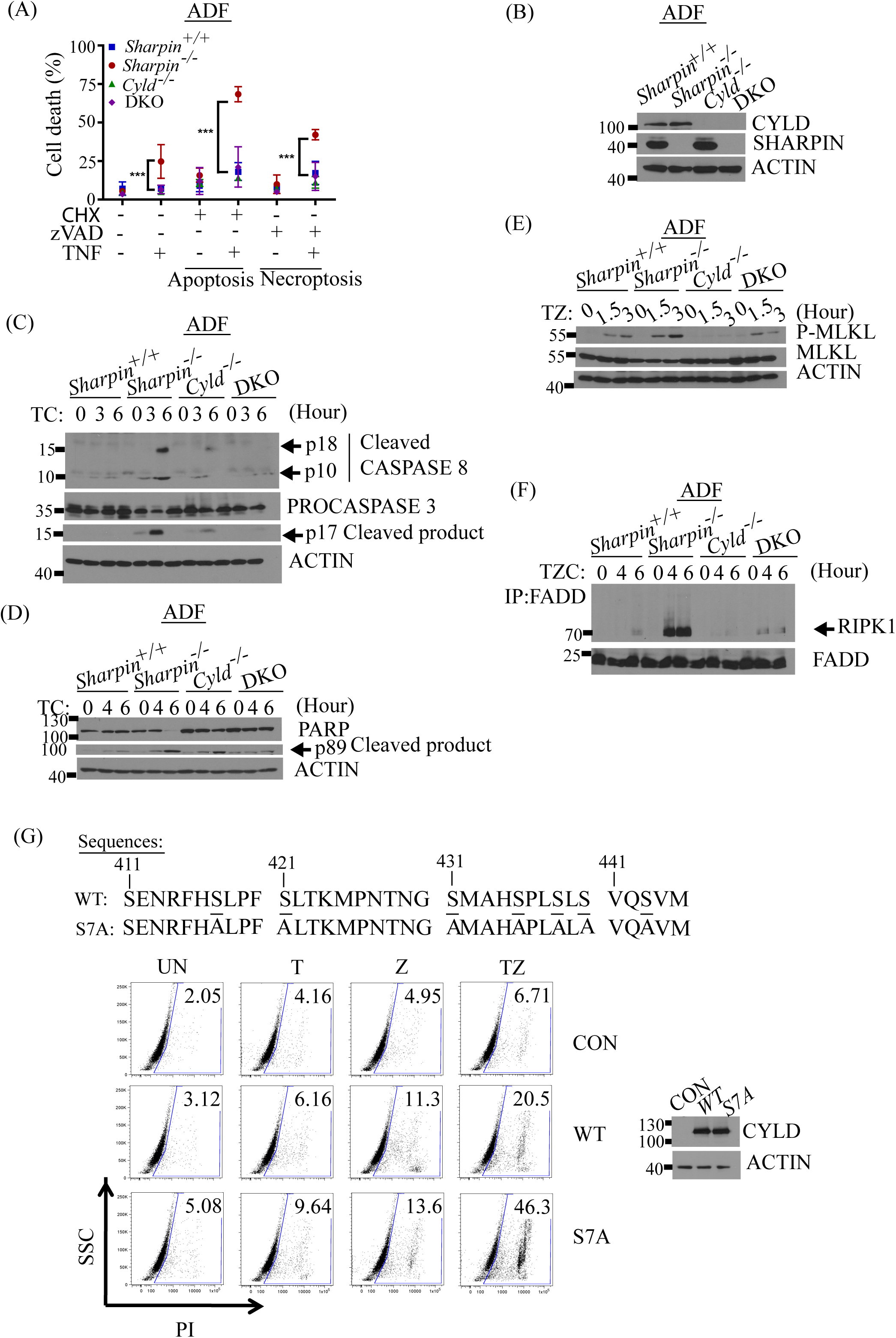
*Sharpin^-/-^* cells are more susceptible to CYLD-mediated cell death. (A) *Sharpin^+/+^*, *Sharpin^-/-^*, *Cyld^-/-^* and DKO ADF were treated with mTNF (10 ng/ml) for 24 h in the presence or absence of zVAD (20 μM) to induce necroptosis or cycloheximide (CHX, 1 μg/ml) to induce apoptosis. Cell death was analyzed by propidium iodide staining and flow cytometry. Data from three independent experiments are shown. Error bars represent mean±SD and one-way ANOVA analysis was performed. ***P<0.001 *Sharpin^-/-^* versus *Sharpin^+/+^* ADF. P value of DKO versus *Sharpin^+/+^* ADF is not significant. (B) CYLD and SHARPIN expression in *Sharpin^+/+^*, *Cyld^-/-^*, *Sharpin^-/-^*, and DKO ADF. (C-D) *Sharpin^+/+^*, *Cyld^-/-^*, *Sharpin^-/-^* and DKO ADF were stimulated with mTNF (100 ng/ml) in the presence of CHX (1 μg/ml) for 3 and 6 h. Apoptosis was examined by blotting for cleaved CASP8 and cleaved CASP3 (C) or cleaved PARP (D). For each panel, data shown are representative of at least three independent experiments. (E) Necroptosis was analyzed by blotting for phospho-MLKL (P-MLKL) in *Sharpin^+/+^*, *Sharpin^-/-^*, *Cyld^-/-^*, and DKO ADF stimulated with mTNF (25 ng/ml) in the presence of zVAD (20 μM) for 1.5 and 3 h. Result shown is representative of at least three independent experiments. (F) *Sharpin^+/+^*, *Sharpin^-/-^*, *Cyld^-/-^*, and DKO ADF were stimulated with mTNF (100 ng/ml) in the presence of CHX (1 μg/ml) and zVAD (20 μM) for 2 and 4 h. The death-inducing signaling complex (DISC) was isolated by FADD immunoprecipitation and sequentially blotted for RIPK1 and FADD. Experiment was repeated twice with similar results. (G) Residues 411 to 445 of wild type mouse CYLD encompassing the cluster of seven serines equivalent to those reported to be phosphorylated on human CYLD, and the corresponding ser-to-ala mutations in the non-phosphorylatable CYLD-S7A mutant are depicted. Ser417 of mouse CYLD is equivalent to Ser418 of human CYLD. *Cyld^-/-^* MEF were transduced with retroviruses encoding a control, wild type CYLD or CYLD-S7A. 24 h after transduction, cells were trypsinized and plated at a density of 3.5×10^4^ cells/well in 24-well plates. The next day, cells were treated with DMSO as negative control (UN), 100 ng/ml mTNF (T), 50 uM zVAD-FMK (Z) or the combination of the two (TZ). Propidium iodide staining and flow cytometry was performed 24 hours after stimulation. Numbers indicate the percentage of PI+ cells. Aliquots of the retroviral transduced MEF were also analyzed by western blotting to demonstrate equivalent expression of wild type and mutant CYLD. Data shown is from one of two independent replicate experiments with similar result.

If *Sharpin^-/-^* cells are more sensitive to death due to defective CYLD phosphorylation, then a non-phosphorylatable CYLD mutant would be predicted to mimic the SHARPIN deficiency (i.e., more sensitive to cell death). Mutations in the cluster of seven serines around residue 418 of human CYLD was shown to result in a gain-of-function in deubiquitinase activity (43). We generated an analogous mutant of mouse CYLD with seven serine to alanine substitutions (CYLD-S7A) and complemented *Cyld^-/-^* MEF with either wild type or mutant CYLD. The complemented cells were treated with TNF in the presence of zVAD-fmk in order to prevent CYLD cleavage by CASP8 (53) and thus negate the potential confounding issue of differential cleavage of CYLD-WT versus CYLD-S7A by CASP8. Under this stimulation condition, CYLD-complemented cells undergo necroptosis (53) and this death response, quantified by PI staining, was enhanced in the CYLD-S7A-complemented cells as compared to CYLD-WT-complemented cells (Figure 4G). The CYLD-S7A-complemented cells phenocopied the SHARPIN-deficient cells (Supplementary Figure S1A & 1B), consistent with the notion that phosphorylation and therefore inhibition of CYLD, is a key regulatory event that suppresses cell death in response to TNF. Our findings imply that there is dysregulated CYLD activity in SHARPIN-deficient cells.

### CYLD-mediated death of myeloid cells causes skin inflammation

The leukocytic infiltration in the skin of the *Sharpin^-/-^* mice consists of granulocytes and macrophages (19) prompting us to examine the role of macrophages in the pathogenesis of skin lesions in these mice. We recently reported that macrophages are highly susceptible to CYLD-dependent auto-necroptosis mediated by the TNF that is produced in response to TLR4 ligation (59). Bone marrow derived macrophages (BMDM) deficient in SHARPIN showed a defect in CYLD phosphorylation when stimulated with TNF (Figure 5A & Supplementary Figure S4A) albeit the defect was modest. A similar defect was also observed when these cells were stimulated with lipopolysacharrides (LPS) to activate TLR4, or polyIC (pIC) to activate TLR3 (Figure 5B & C, Supplementary Figure S4B & C). Therefore, we tested to see if these macrophages also exhibit altered necroptosis. In response to low-dose LPS that elicits TNF-dependent necroptosis (59), *Sharpin^-/-^* BMDM were more sensitive to necroptosis compared to their wild type counterparts and this was abrogated in the *Sharpin^-/-^Cyld^-/-^* BMDM (Figure 5D). A similar effect was also observed using pIC to trigger necroptosis (Figure 5E). Analysis of DISC formation showed that RIPK1 was detected in FADD pulldowns of *Sharpin^-/-^* BMDM more rapidly than in *Sharpin^+/+^* cells treated with TNF (Figure 5F). Similar to that observed previously in ADF (Figure 4F), RIPK1 association with the DISC was abrogated in *Sharpin^-/-^Cyld^-/-^* BMDM. To confirm the *in vitro* BMDM data, we looked for evidence of macrophage death in skin sections taken from mice of the different genotypes. Consistent with a previous report showing apoptosis occurring in the skin of *Sharpin^-/-^* mice (60), we detected cleaved CASP3 in the skin of *Sharpin^-/-^* mice, a proportion of which co-localized with F4/80+ macrophages (Figure 5G & H). However, cleaved CASP3 was largely absent in the *Sharpin^-/-^Cyld^-/-^* skin consistent with our *in vitro* data showing that SHARPIN-deficient cells undergo cell death in a CYLD-dependent manner.

**Figure 5.**
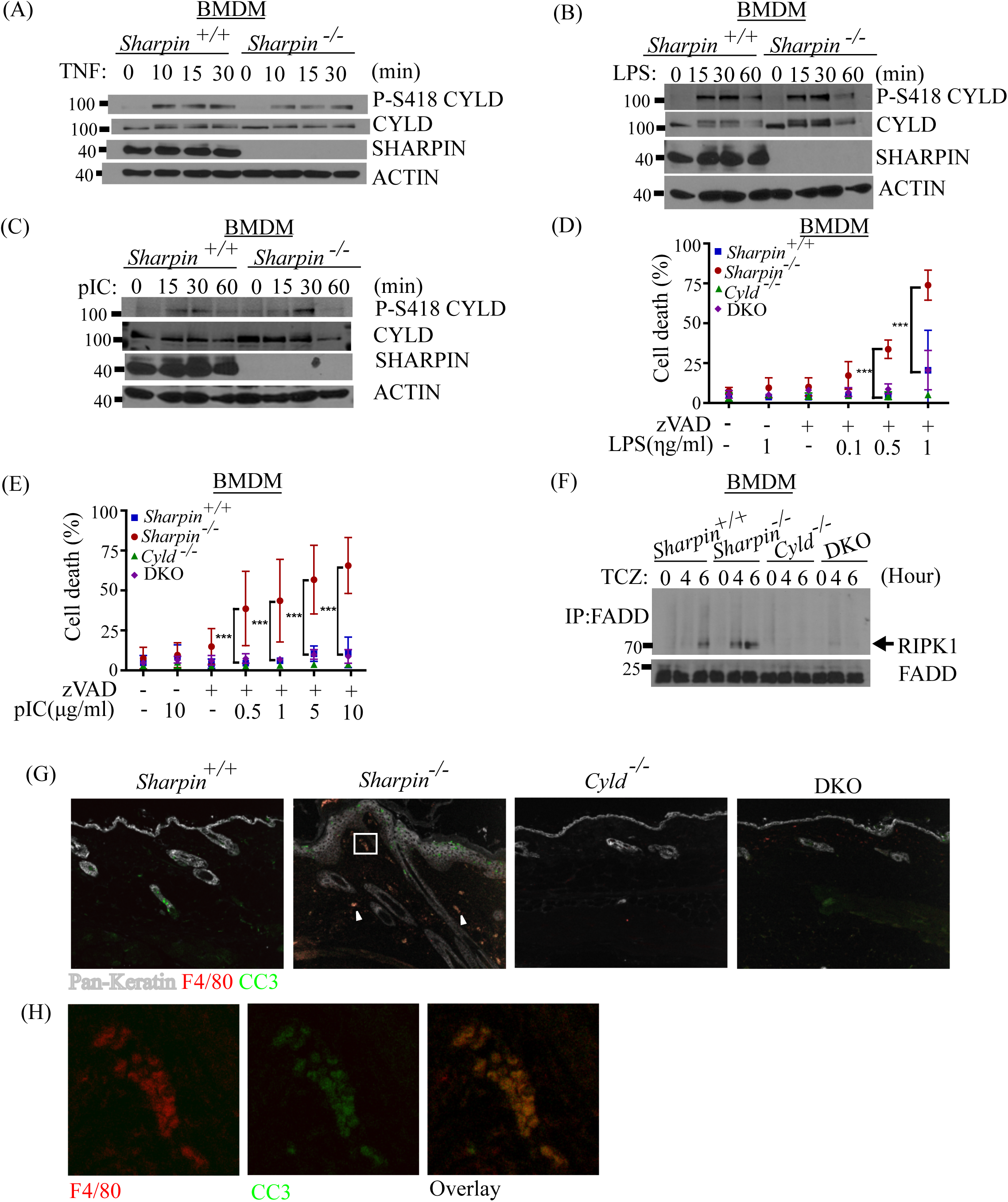
*Sharpin^-/-^* myeloid cells are more susceptible to CYLD-mediated cell death. (A-C) *Sharpin^+/+^* and *Sharpin^-/-^* bone marrow derived macrophages (BMDM) were stimulated with (A) 100 ng/ml mTNF, (B)10 ng/ml LPS and (C) 2.5 μg/ml polyIC for the indicated times. CYLD phosphorylation was analyzed as in Figure 3A. Blots shown are representative of three independent experiments performed with each stimulus. (D-E) *Sharpin^+/+^*, *Sharpin^-/-^*, *Cyld^-/-^* and DKO BMDM were stimulated with the indicated doses of (D) LPS and (E) polyIC in the presence of DMSO or zVAD (20 μM) for 24 h. Cell death was analyzed as in Figure 4A. Error bars represent mean±SD and one-way ANOVA analysis was performed. ***P<0.001 *Sharpin^-/-^* versus *Sharpin^+/+^* BMDM. P value of DKO versus *Sharpin^+/+^* BMDM is not significant. (F) *Sharpin^+/+^*, *Sharpin^-/-^*, *Cyld^-/-^* and DKO BMDM were stimulated with mTNF (100 ng/ml) in the presence of CHX (0.25 μg/ml) and zVAD (20 μM) for 4 and 6 h. Translocation of RIPK1 to the DISC was analyzed as in Figure 4F. Result shown is representative of three independent experiments performed. (G) Skin sections from mice of the indicated genotypes were labeled by immunofluorescence with anti-F4/80 and anti-cleaved CASP3 (CC3) to detect macrophages and dying cells, respectively. Representative photomicrographs (100X) from each genotype are shown. White arrowheads indicate co-localization of macrophages with cleaved CASP3. (H) Rectangular insert of the *Sharpin^-/-^* skin section from panel (G) above is enlarged for visualization of the F4/80 and cleaved CASP3 signals.

The genetic crosses conducted using germline knockouts showed CYLD to be the cause of the phenotype observed in *Sharpin^-/-^* mice. However, the contribution of CYLD-dependent death in specific cellular compartments to the *Sharpin^-/-^* phenotype is not clear. The data in Figure 5A-H prompted us to ask whether CYLD-dependent macrophage death has any role in the pathogenesis of the *Sharpin^-/-^* mice. We crossed the *Sharpin^-/-^* strain to the *LysM-cre*-driven conditional knockout of *Cyld* (*Cyld^M-KO^*) we previously generated (59). The *Sharpin^-/-^Cyld^M-KO^* mice showed significant improvement in their dermatitis compared to the *Sharpin^-/-^* mice, which develop discernable dermatitis by week 8 in our colony (Figure 6A). In our cohort, two thirds of *Sharpin^-/-^Cyld^M-KO^* mice did not show skin inflammation by week 16 while the remaining 1/3 had delayed progression (Supplementary Figure S4D). Histological analysis showed that the typical ulceration, thickening of the epidermis and leukocytic infiltration observed in *Sharpin^-/-^* skin was absent in *Sharpin^-/-^Cyld^M-KO^* mice with no disease (Figure 6B). However, other defects in the spleen and thymus of *Sharpin^-/-^* mice were not reversed in the *Sharpin^-/-^Cyld^M-KO^* mice (Figure 6C & D). We confirmed that BMDM generated from *Sharpin^-/-^Cyld^M-KO^* mice were more resistant to cell death than those from *Sharpin^-/-^* mice and that the *LysM-cre* was effective in knocking out CYLD in our BMDM cultures (Supplementary Figure S4E-G). We also crossed the *Sharpin^-/-^* strain to a *Cx3cr1-cre*-driven conditional knockout of *Cyld* (*Cyld^MP-KO^*), which targets mononuclear phagocytes including macrophages (61). The *Cx3cr1-cre*-mediated deletion of *Cyld* recapitulated the effect observed earlier with *LysM-cre* (Supplementary Figure S5A-G). Results from the use of two different *cre* strains that can delete *Cyld* in macrophages led us to conclude that CYLD-dependent macrophage cell death may be a major driver of the inflammatory response in the skin of the *Sharpin^-/-^* mice.

**Figure 6.**
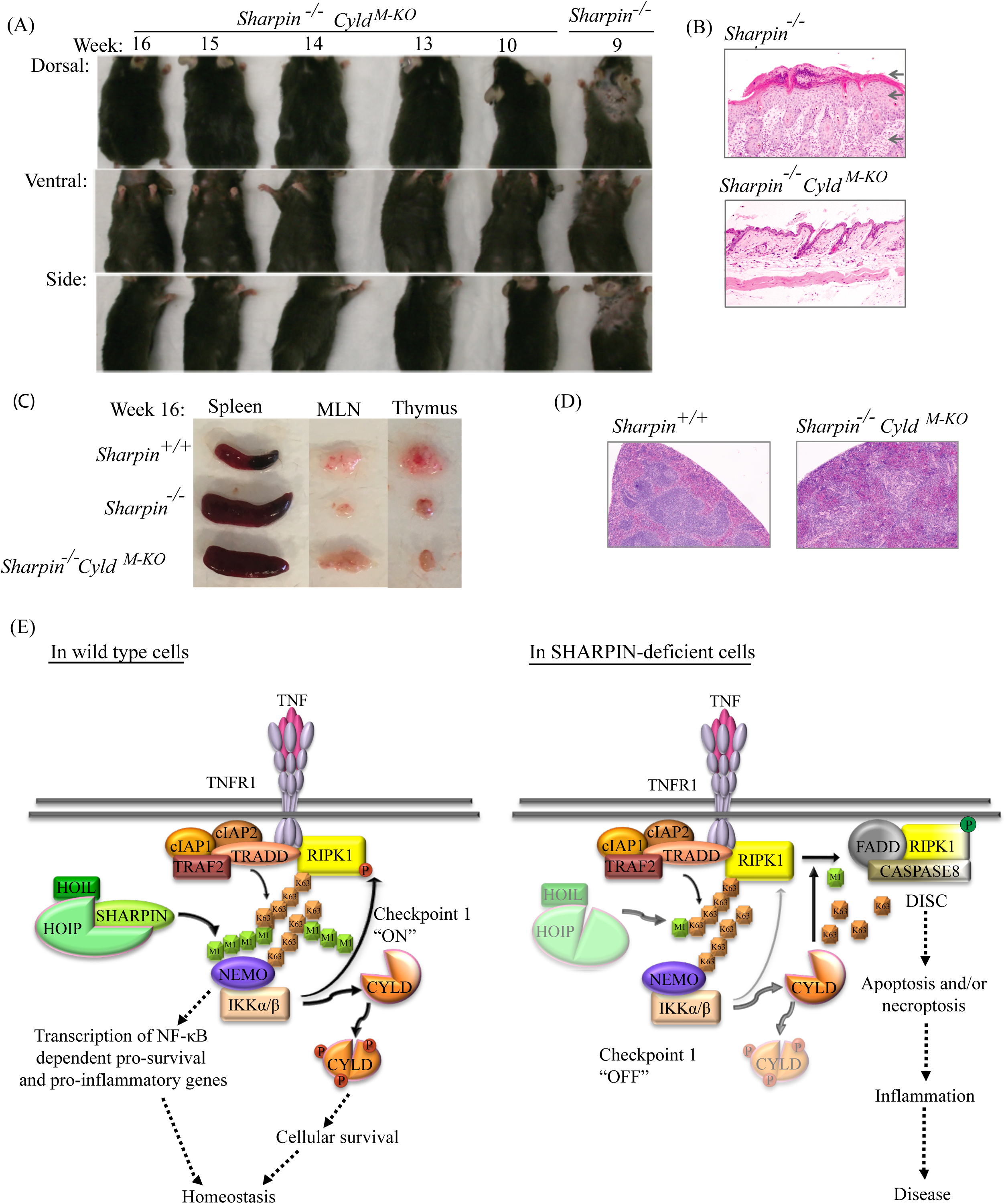
CYLD-mediated death of myeloid cells is necessary to cause skin inflammation in *Sharpin^-/-^* mice. (A) Photograph of 10-16-week-old *Sharpin^-/-^Cyld^M-KO^* mice in comparison to a 9-week-old *Sharpin^-/-^* mouse. (B) H&E staining of skin sections from 16-week-old *Sharpin^-/-^ and Sharpin^-/-^Cyld^M-KO^* mice (100X). Top arrow: orthokeratotic hyperkeratosis and serocellular crust; middle arrow: thickened epidermis; bottom arrow: thickened dermis. (C) Gross analysis of spleens, mesenteric lymph nodes and thymi from 16-week-old *Sharpin^+/+^, Sharpin^-/-^ and Sharpin^-/-^Cyld^M-KO^* mice. (D) H&E staining of spleen sections from 16-week-old *Sharpin^+/+^ and Sharpin^-/-^Cyld^M-KO^* mice. (E) Model of CYLD suppression by linear ubiquitin. In wild type cells, the LUBAC-dependent IKK phosphorylates CYLD to inactivate it. LUBAC could also suppress CYLD via a phosphorylation-independent mechanism. These mechanisms suppress CYLD thereby keeping RIPK1 in a survival mode. In addition, IKK can directly phosphorylate RIPK1 to inhibit its death-signaling function. Linear ubiquitin cooperates positively with K63-linked ubiquitin to maintain cell survival by preventing the dismantling of K63-linked ubiquitin chains. In SHARPIN-deficient cells, there is diminished suppression of CYLD. Therefore, CYLD is active and removes K63-linked ubiquitin chains from RIPK1. This enables RIPK1 to associate with the DISC to induce cell death leading to multi-organ inflammation, abnormal lymphoid tissues and immunodeficiency.

## Discussion

This study now provides genetic data that could account for the *Sharpin^-/-^* phenotype. We propose that in wild type cells, SHARPIN-dependent regulation of CYLD suppresses its function, allowing K63-linked ubiquitin chains on RIPK1 to be sustained thus preventing RIPK1 from becoming a death-signaling molecule (Figure 6E, left panel). Thus, SHARPIN-dependent inhibition of CYLD is an essential component of Checkpoint 1 in the TNF signaling pathway that keeps cell death at bay. In the absence of SHARPIN, there is a failure to suppress CYLD, which dismantles K63-linked ubiquitin chains from RIPK1 to initiate the RIPK1-dependent death cascade (Figure 6E, right panel). We note that there is reduced phosphorylation of serine 418 of CYLD in SHARPIN-deficient cells, which is known to be phosphorylated by IKKβ and IKKε (43, 57). Reduced CYLD phosphorylation could be due to reduced IKK activity in SHARPIN-deficient cells (7–9). In addition, CYLD is recruited to the receptor complex in a SPATA2 and RNF31/HOIP–dependent manner (44–48) and since *Sharpin^-/-^* cells have reduced RNF31/HOIP level (7–9), this could reduce CYLD recruitment to the receptor complex and subsequent phosphorylation by IKK. In light of published studies that phosphorylation of CYLD at a cluster of serines around residue 418 inhibits its enzymatic activity (43, 57), it is likely that the defective phosphorylation of CYLD in the SHARPIN-deficient cells enhanced CYLD’s removal of ubiquitin chains from RIPK1. Alternatively, phosphorylation could regulate CYLD function in a non-enzymatic manner such as regulating its localization or its interaction with signaling partners. Furthermore, it is possible that SHARPIN could inhibit CYLD through some other mechanism independent of phosphorylation and this will be further explored.

In addition to phosphorylation, CYLD can also be negatively regulated by proteolysis mediated by CASP8 and MALT1 (53, 56). Interestingly, SHARPIN-deficient splenocytes may circumvent the defect in CYLD phosphorylation by upregulating these proteolytic events suggesting that these inhibitory mechanisms could compensate for each other. Furthermore, CYLD can be phosphorylated by multiple members of the IKK family including IKKβ, IKKε and TBK1 suggesting that there may be redundancy in regulating CYLD phosphorylation in different cell types. This redundancy could explain the modest defect in CYLD phosphorylation in the *Sharpin^-/-^* BMDM as macrophages express IKKεand TBK1 (62). Besides CYLD, these IKK kinases also directly phosphorylate RIPK1 to inhibit the death-signaling function of RIPK1 (26–30). Taken together, the emerging picture is that a fully functional Checkpoint 1 needed to suppress ripoptocide is dependent on multiple post-translational modification of several molecules.

Biologically, these results really point towards suppression of CYLD as central to immune homeostasis as perturbation in CYLD regulation, as observed in the *Sharpin^-/-^* mice, leads to multi-organ inflammation and disruption of lymphoid tissues. It also demonstrates that CYLD functions as a pro-inflammatory molecule since its deletion resolves the inflammation seen in the *Sharpin^-/-^* mice. These aspects of CYLD in the proper functioning of the immune system have not been appreciated to date. Previous studies with CYLD have described it as having an anti-inflammatory function (63–65), primarily because of its inhibitory effect on the NF-κB pathway. We now show that CYLD can be highly pro-inflammatory via its death-inducing function, which is revealed upon the loss of an upstream inhibitory mechanism (e.g., SHARPIN/LUBAC). This is a significant advance in our understanding of the role played by ubiquitin-modifiers in regulating immune homeostasis and response. Our studies strongly suggest that the defects observed in human patients with mutations in *RNF31/HOIP* and *RBCK1/HOIL1* (17, 18) could be due to inappropriate CYLD activity.

Excessive CYLD-dependent cell death in the *Sharpin^-/-^* mice can cause skin inflammation due to (1) death of epithelial cells that could result in cellular erosion and ulcers in and of itself, (2) loss of barrier function in the epithelial layer and subsequent breach by commensal microbiota and environmental contaminants (bedding and food), and/or (3) excessive cellular debris from dying cells that are not properly cleared by phagocytes. The reversal of skin inflammation in *Sharpin^-/-^* mice following deletion of *Fadd* or *Tradd* in keratinocytes indicates that cell death in this cellular compartment is critical for inflammation (22). Our observation that the skin inflammation was also reduced by *LysM-cre*- and *Cx3cr1-cre*-mediated deletion of *Cyld* suggested that cell death of macrophages also has a pro-inflammatory function. One explanation is that death of *Sharpin^-/-^* epithelial cells is the initiating factor for the skin inflammation while death in macrophages plays an accessory role. SHARPIN-deficient macrophages can contribute to the skin inflammation by undergoing more death in response to invading microbiota and thereby amplifying the inflammatory response. Alternatively, because SHARPIN-deficient macrophages are more sensitive to death, they may be less effective in clearing up debris from dying epithelial cells. Since both *LysM-cre*- and *Cx3cr1-cre*-mediated deletion of *Cyld* had the same effect, our data strongly suggest that improper cell death in macrophages is highly inflammatory in the skin but does not have a role in other immune dysfunction in the *Sharpin^-/-^* mice such as the disruption in splenic architecture.

The *Sharpin^-/-^* mouse illustrates the pathophysiology when the TNF response switches from cell survival to cell death by virtue of a genetic defect in CYLD regulation. The question remains as to the physiological function of this molecular switch in a normal individual and the role of TNF-mediated cell death. Indeed, this question has remained unresolved for over forty years despite the fact that this was the first cellular response attributed to TNF, as reflected in the name of this cytokine (66). This conserved cell-killing function of TNF has been postulated to serve an evolutionary role against microbial infection (38, 67). The analysis of the *Sharpin^-/-^* mice suggests that a biological consequence of TNF/CYLD-mediated ripoptocide is inflammation. In particular, the invocation of this death response in macrophages is highly inflammatory since blocking this from occurring in macrophages ameliorated the skin inflammation of the *Sharpin^-/-^* mice. Therefore, it is probable that the inflammation caused by ripoptocide has an anti-microbial function, as previously postulated (67). Since CYLD’s death-inducing function can be controlled by post-translational modifications, the speculation is that this early transcription-independent molecular switch may be disrupted by pathogens, thus providing a facile and rapid switch to cell death to generate an inflammatory response beneficial to the host. The IKK complex that phosphorylates CYLD is also necessary for induction of NF-κB-dependent inflammatory genes. Therefore, pathogens may encode inhibitors to block IKK as a strategy to limit the host expression of inflammatory genes and in so doing would also impair CYLD phosphorylation. This would be expected to unleash CYLD-dependent ripoptocide and the ensuing inflammation would function as a countermeasure against blockade of inflammatory gene induction by pathogens. It was particularly interesting that blocking CYLD-dependent ripoptocide in macrophages had a strong anti-inflammatory effect on the dermatitis of the *Sharpin^-/-^* mice. These sentinel cells are the major producers of inflammatory cytokines in response to infection and are also often the host for replicative parasitic pathogens. Therefore, the presence of this CYLD-regulated ‘trap door’ in macrophages would be advantageous to the host. An example of this is infection by the bacteria pathogen *Yersinia*. It encodes an effector molecule YopJ to block cytokine gene synthesis, which likely leads to the disruption of Checkpoint 1 to trigger ripoptocide and this has been shown to be beneficial to the host (29, 68). It is likely that there are other pathogens that behave similarly.

In summary, we have uncovered the mechanistic basis for the immune dysfunction observed in linear ubiquitin-deficient mice. This involves a defect in regulating CYLD, a molecular switch that is normally turned off to prevent TNF from inducing death. The finding that linear ubiquitin regulates the function of CYLD, an enzyme that dismantles K63-linked ubiquitin chains, reveals positive cooperativity between the two forms of ubiquitin modifications in regulating a cellular response. In addition to providing an explanation for the immune system abnormalities caused by genetic defects in linear ubiquitination, this molecular insight significantly advances our understanding of how inflammation and immunodeficiency may be regulated by cell death.

## Materials & Methods

### Reagents

Mouse and human recombinant TNF (Peprotech), zVAD-fmk (Calbiochem and Bachem), necrostatin-1 (Tocris Bioscience), cycloheximide (Sigma), TPCA-1 ([5-(*p*-Fluorophenyl)-2-ureido]thiophene-3-carboxamide, EMD Millipore), LPS *E.coli* 0111:B4 (Sigma) and poly(I:C) HMW (Invivogen) were obtained from the indicated sources. RBC lysis buffer (Invitrogen) and Protease Inhibitor Cocktail Set V (EMD Millipore) were purchased from the indicated sources. Lentiviral constructs encoding a non-targeting shRNA or CYLD-targeting shRNA (SHCLNG-NM_173369) were obtained from Sigma.

### Antibodies

Antibodies were purchased from the indicated vendors. GAPDH clone D-6 (SC166545), FADD clone M19 (SC6063), LAMININ clone C-20 (SC6216), TUBULIN clone B-7 (SC5286), NF-κB p65 clone F-6 (SC8008), and goat IgG HRP (SC2350) from Santa Cruz; CYLD clone D1A10 (#8462S), P-S418-CYLD (#4500S), CASPASE 8 (#4927S), cleaved CASPASE 8 (#9496S), CASPASE 3 (#9662S), cleaved CASPASE 3 (#9661S), PARP (#9532S), cleaved PARP (#9541S), β-ACTIN (#3700S), and mouse IgG HRP (#7076S) from Cell Signaling; F4/80 (#14-4801-82) from eBioscience; Pan-Keratin (#ab8068) and CD45 (#ab10558) from Abcam; RIPK1 (#610458) from BD Transduction Laboratories; RIPK3 (#2283, ProSci); P-MLK clone 7C6.1 (#MABC1158) and MLKL clone 3H1 (#MABC604) from EMD Millipore; SHARPIN (#14626-1-AP) from Proteintech; Rabbit Ig HRP (#111-035-144) from Jackson Immuno Research Labs.

### Mice

*Sharpin^-/-^Cyld^-/-^* mice were generated by crossing *Sharpin^-/-^* (C57BL/KaLawRij-*Sharpin^cpdm^*/RijSunJ) provided by Dr. John Sundberg with *Cyld^-/-^* mice (B6;129S-*Cyld^tm1Scs^*) provided by Dr. Shao-Cong Sun (69). Experiments were conducted using *Sharpin^+/+^*, *Sharpin^-/-^* and *Cyld^-/-^* littermates as controls. *Sharpin^-/-^Cyld^M-KO^* mice were generated by crossing *Sharpin^-/-^* strain with the *Cyld^flox/flox^* (B6.129S-*Cyld^tm1.1Attg^*) *x LysM-cre* (B6.129P2-*Lyz2^tm1(cre)Ifo^*/J) strain previously described (59). *Sharpin^-/-^Cyld^MP-KO^* mice were generated by crossing *Sharpin^-/-^* strain with the *Cyld^flox/flox^* (B6.129S-*Cyld^tm1.1Attg^*) *x Cx3cr1-cre* (B6J.B6N(Cg) *Cx3cr1cre^tm1.(cre)Jun^*/J) strain. The *Cx3cr1-cre* was provided by Dr. Sergio Lira. All experiments involving the use of animals were performed in agreement with approved protocols by the Institutional Animal Care and Use Committee (IACUC) at the Icahn School of Medicine at Mount Sinai.

### Histology

Skins, spleens, colons and small intestines were removed, fixed in 10% neutral-buffered formalin, and then transferred into 70% ethanol. Paraffin embedding, tissue sectioning, H&E staining and immunohistochemistry were performed by the Comparative Pathology Lab in the Center for Comparative Medicine and Surgery, or by the Biorepository and Pathology CoRE at the Icahn School of Medicine at Mount Sinai. Immunofluoresence labeling of skin sections was carried using the protocol previously described (70).

### Clinical scoring

The gross scoring scheme was modified from a system developed for a different mouse disease, ulcerative dermatitis that occurs in C57BL/6 substrains (71). The clinical score combines gross evaluation with histologic scoring from skin sections. In brief, 4 criteria were evaluated: character of lesion (0=none, 1=alopecia or excoriations only, or 1 small punctuate crust, 2=multiple, small punctuate crusts or coalescing crusts (>2mm), regions affected (head/cervical, thoracic, and/or abdominal/caudal; 0=none, 1=<25%, 2=25-50%, 3=>50%). The “regions affected” score in Hampton et al (71) was based on the progression of the disease. Ulcerative dermatitis in B6 mice often started in the intercapsular/dorsal back area (region 2) and progressed, thus encroachment on the head and face (region 1) was considered to be a severe score. In this study, the lesions often start in the ventral chin/neck region (region 1) and progress to the thorax, abdomen and back. The scoring system was modified to increase in severity based on the number of regions affected rather than the specific region affected because of this difference in progression of skin disease. For the histologic lesions, scoring was a subjective 0=none, 1=mild, 2=moderate, 3=marked scale. The scoring of inflammation in the dermatitis was done using the following scale 0=none, 1=mild (focal or few foci of inflammation in superficial dermis), 2=moderate (multiple foci with inflammation in deep dermis, with or without ulcerations), and 3=severe (regionally extensive with inflammation extending to the subcutis and ulceration).

### Serum immunoglobulins, cytokines and chemokines assessment

Blood was collected via facial vein from 10-16 weeks old mice and spun at 1000 x *g* for 10 min at room temperature. Serum was then collected and stored at −80°C. IgA, IgE, IgG and IgM in the serum were quantified using multiplex Luminex*®* Immunoglobulin Isotyping. IFNG, IL2, IL5, IL6, IL12/23p40, eotaxin and CCL3 (MIP-1α) in the serum were quantified using Luminex*®*Cytokines and Chemokines Multiplex Assays.

### Stool IgA assessment

Stool pellets obtained from 10-16 week old mice were placed in chilled PBS containing protease inhibitors (100 mg/ml), repeatedly smashed and vortexed for 4-5 times. The suspension was then centrifuged at 8000 x *g* for 10 min at 4 °C. The supernatants were collected and frozen at −80°C. IgA levels were quantified using the Mouse IgA ELISA Quantitation Set (Bethyl Laboratories).

### Primary adult dermal fibroblast (ADF) cultures

A small section (2 mm x 3 mm) of the ears from adult mice were snipped, soaked in 70% ethanol for 30 sec and washed with sterile PBS. Tissues were then diced in culture media (DMEM with 10% FBS and antibiotics) and spun down. The pellets were trypsinized for an hour at 37 °C with intermittent hard vortexing prior to centrifugation. The pellets were then plated onto a tissue culture dish and cultured in DMEM with 10% FBS and antibiotics at 37°C with 5% CO_2_ for 1-2 weeks.

### Total splenocytes and B cell isolation

Spleens from 8-12-week-old adult mice were mashed and filtered through 70 μM sterile funnel and with RBC lysis buffer for 5 min. Total splenocytes were then rested at 37 °C in a 5% CO_2_ incubator for at least 2h before use. For B cell preparation, total splenocytes were subjected to negative selection using BioLegend MojoSort Magnetic Cell Separation kit to obtain purified B cells. The purity was determined by labeling for CD19 or CD40 by flow cytometry (>90%). B cells were rested for at least 2h before use.

### Bone marrow-derived macrophage cultures

Bone marrow-derived macrophages (BMDM) were generated by culturing bone marrow progenitors from femurs and tibias of mice in RPMI containing 10% FBS and 30% L929 conditioned medium as described (72). Day 8 to 11 cultures were used for all experiments.

### Retroviral transduction of MEF

HEK293 EBNA cells were transfected by calcium phosphate precipitation with plasmids encoding VSV-G and GAG-POL, together with an MMLV-based retroviral expression construct encoding control or relevant genes. 48 h post-transfection, the viral supernatants were collected and used to infect MEF by spinoculation. MEF were selected with puromycin or blasticidin 24 h subsequently. *Sharpin^-/-^* MEF were complemented with a negative control or wild type mouse *Sharpin* gene. Both lines were asecondarily transduced with control or IκBSR to block NF-κB signaling. *Sharpin^-/-^Cyld^-/-^* MEF were complemented with (i) control gene, (ii) mouse *Sharpin* wild type alone, (iii) mouse *Cyld* wild type alone, or (iv) both mouse *Sharpin* and *Cyld* wild type alleles. *Cyld^-/-^* MEF were complemented with (i) control gene, (ii) mouse *Cyld* wild type allele, or (iii) mouse *Cyld^S7A^* mutant allele. The *Cyld^S7A^* ser-to-ala mutations were generated using Quikchange Multi Site-Directed Mutagenesis Kit (Agilent).

### CYLD knockdown in MEF

HEK 293 EBNA cells were transfected by calcium phosphate precipitation with plasmids encoding VSV-G and GAG-POL, together with Mission*®* shRNA lentiviral vectors targeting CYLD or a non-targeting shRNA. Lentiviral infection of *Sharpin^-/-^* MEF was carried out as described above for retroviral transduction.

### Cell death assays

Cellular death analyses were performed by Annexin V or propidium iodide staining and flow cytometry as previously described (53). Cell death was also quantified using the CellTiter-Glo Luminescent Cell Viability Assay (Promega). Biological triplicates were used in the experiments for ADF and BMDM cell death analysis.

### Co-immunoprecipitation

For FADD immunoprecipitation, 2 x 10^6^ cells ADF and MEF or 3-4 x 10^6^ BMDM per sample were stimulated and then lysed in buffer containing 20 mM Tris pH 7.4, 150 mM sodium chloride, 10% glycerol, 0.2% NP-40, 0.1 mM sodium orthovandate, 5 mM β-glycerophosphate and Protease Inhibitor Cocktail Set V for 20 min on ice. Lysates were cleared by centrifugation at 10, 000 x *g* at 4°C and protein concentration measured using Pierce BCA (Thermo Fisher). Equivalent amount of protein in each sample were immunoprecipitated by rotating with 0.25-1 µg of FADD antibody (M19) overnight at 4°C. Immune complexes were precipitated with Protein A/G beads. After extensive washing, the beads were eluted with SDS-sample buffer at 70°C for 20 min. Blotting with anti-RIPK1 was carried out subsequently.

For RIPK1 immunoprecipitation, 0.84 x 10^6^ MEF stimulated and then lysed in buffer containing 20 mM Tris pH 7.4, 150 mM sodium chloride, 10% glycerol, 1% TritonX-100, 0.5mM DTT, 10mM N-Ethylmaleimide, 0.1 mM sodium orthovandate, 5 mM β-glycerophosphate and Protease Inhibitor Cocktail Set V for 20 min on ice. Lysates were cleared by centrifugation at 10, 000 x *g* at 4°C. One tenth of the lysate was set aside for SDS-PAGE and sequential blotting with anti-RIPK1 and anti-ACTIN. The reminder of the lysates was immunoprecipitated by rotating with 0.4 µg of RIPK1 antibody overnight at 4°C. Immune complexes were precipitated with Protein A/G beads. After extensive washing, the beads were eluted with SDS-sample buffer at 70°C for 20 min. Blotting with anti-RIPK1 was carried out subsequently.

### Western blot Analysis

Cellular lysates obtained using 1% Triton X-100 were resolved by reducing SDS PAGE. For P-MLKL, lysates were generated using M2 buffer as previously described (73). Blotting was conducted using standard techniques and detected by LiCor Odyssey or by chemiluminescence. Semi-quantification of the blots was carried using Image J software (74).

### Statistical Analysis

Statistics were performed using one-way Anova, Student’s t-test or Mann Whitney U test. (*p<0.05, **p<0.01, ***p<0.001)

## Author contributions

R.L.A designed and performed the experiments. J.P.S. and S.C.C. provided *Sharpin^-/-^* and *Cyld^-/-^* mice, respectively. V.G. did the clinical scoring in the double-blind study. J.P.S. reviewed the histopathology. H.X. and S.A.L. provided technical guidance. P.S.H. provided critical comments, edited the manuscript and provided financial support through fellowship grants. R.L.A. and A.T.T. wrote the manuscript and directed the studies.

## Acknowledgements

We would like to thank the following from the Icahn School of Medicine at Mount Sinai: Mr. Alan Soto from Biorepository and Pathology CoRE services for processing the histology, Ms. Ying Dai from the Comparative Pathology Laboratory in the Center for Comparative Medicine and Surgery for imaging, and Drs. Thomas Krauss and Thomas Moran for the use of the Luminex instrument. This work was supported by National Institutes of Health (NIH) grants AI052417 (A.T.T.), AI104521 (A.T.T.), AI132405 (A.T.T & P.S.H.), DK072201 (S.A.L. & A.T.T.), CA161373 (S.A.L.), AI064639 (S.-C. S) and AR049288 (J.P.S.). This work was also supported by a Senior Research Award #253097 (A.T.T.) and #330239 (S.A.L.) from the Crohn’s and Colitis Foundation of America. R.L.A was supported by National Institute of Health training grants: AI078892, GM062754, and A1007605. We declare that there are no financial conflicts of interest.

