## Supplemental Data for "Immune dysregulation in SHARPIN-deficient mice is dependent on CYLD-mediated cell death"

Supplementary Figure S1

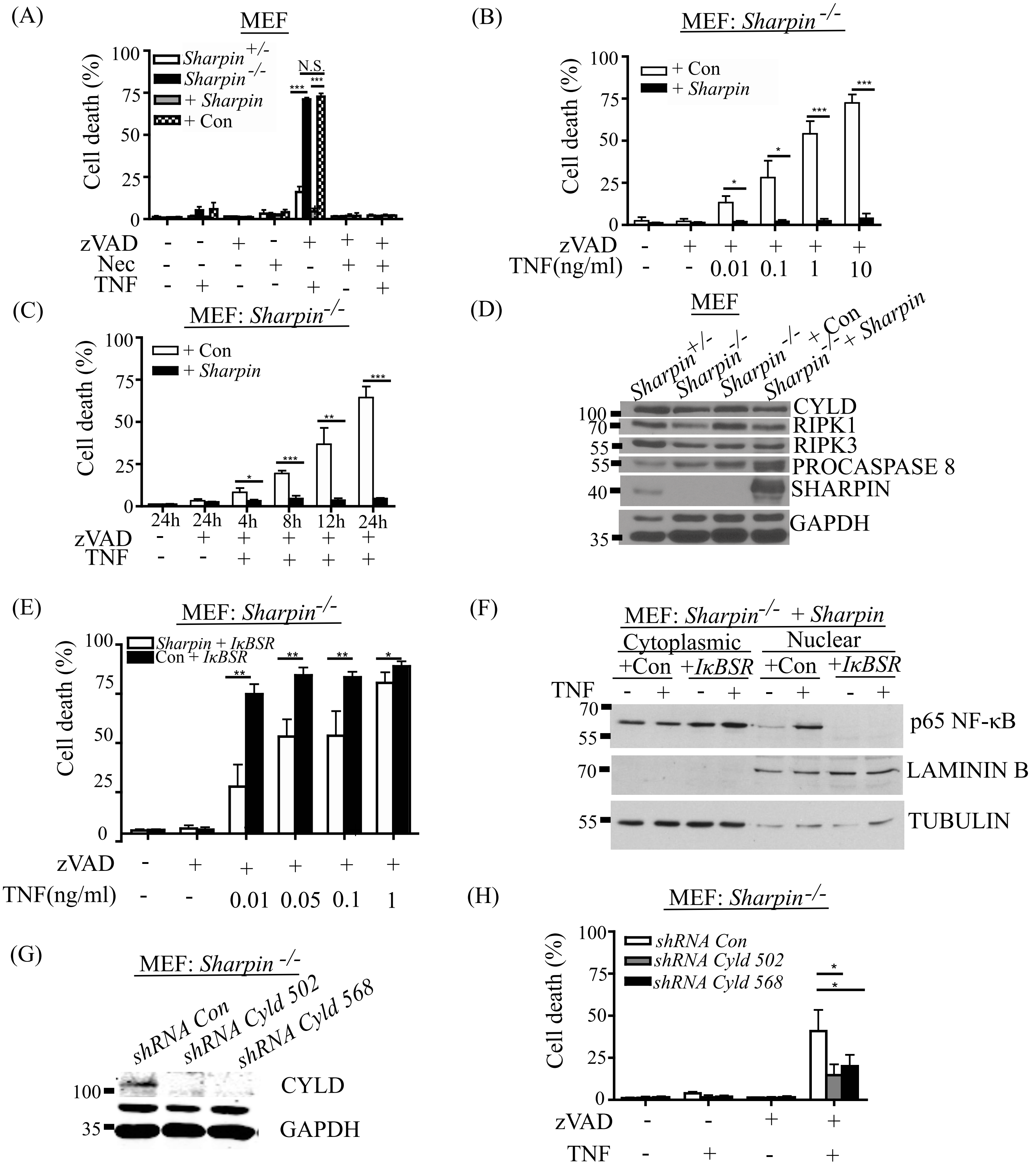

**Supplementary Figure S1. SHARPIN-deficient MEF are sensitized to CYLD-dependent cell death.**

(A) Mouse embryonic fibroblasts (MEF) derived from *Sharpin*$^{-/-}$ mice were first complemented with a retroviral vector encoding *Sharpin* (indicated by +*Sharpin*) or bacterial GST as a control gene (indicated by +*Con*). These two lines together with *Sharpin*$^{+/-}$ and *Sharpin*$^{-/-}$ MEF were analyzed for sensitivity to TNF-induced necroptosis by Annexin V staining. The four lines were stimulated with hTNF (10 ng/ml) in the presence or absence of zVAD (50 μM), Necrostatin (30 μM) or both for 24 h. The results confirmed that the *Sharpin*$^{-/-}$ cells were more sensitive to TNF-induced necroptosis and this was reversed when these cells were complemented with *Sharpin* but not a control gene.

(B) *Sharpin*$^{-/-}$+*Sharpin* and *Sharpin*$^{-/-}$+*Con* MEF were stimulated with increasing dose of hTNF in the presence of zVAD (50 μM) for 24h followed by Annexin V staining. SHARPIN-deficient cells were more sensitive to TNF-induced necroptosis in a dose-dependent manner.

(C) *Sharpin*$^{-/-}$+*Sharpin* and *Sharpin*$^{-/-}$+*Con* MEF were stimulated with hTNF (10 ng/ml) in the presence of zVAD (50 μM) for different times followed by Annexin V staining. SHARPIN-deficient cells were more sensitive to TNF-induced necroptosis in a time-dependent manner.

(D) Protein expression of CYLD, RIPK1, RIPK3, PROCASP8 and SHARPIN in *Sharpin*$^{+/-}$, *Sharpin*$^{-/-}$, *Sharpin*$^{-/-}$+*Sharpin* and *Sharpin*$^{-/-}$+*Con* MEF.

(E) *Sharpin*$^{-/-}$+*Sharpin* and *Sharpin*$^{-/-}$+*Con* MEF were stably transfected with the non-degradable IκBα super-repressor (IκBSR) to block NF-κB. The two cell lines were treated with increasing doses of hTNF in the presence of zVAD (50 μM) for 24 h prior to Annexin V staining. In the absence of NF-κB signaling, the SHARPIN-deficient cells remained more sensitive to TNF-induced necroptosis demonstrating a pro-survival role for SHARPIN independent of NF-κB.

(F) Cytoplasmic and nuclear extracts were prepared from *Sharpin*-reconstituted MEF stably transfected with either control or the IκBSR after hTNF stimulation. Extracts were blotted with an antibody against NF-κB p65/RELA, which showed a stimulus-dependent translocation of p65 to the nucleus in control-transfected cells but this was blocked in the IκBSR-transfected cells. This result demonstrates the effectiveness of the IκBSR in blocking NF-κB in the MEF used in (E) above.

(G) *Sharpin*$^{-/-}$ MEF with stable transfection of a non-targeting hairpin (*shRNA Con*) or two different hairpins targeting CYLD (*shRNA Cyld 502* and *shRNA Cyld 568*) were blotted with anti-CYLD to confirm knockdown of CYLD.

(H) *Sharpin*$^{-/-}$ MEF with control or CYLD knockdown from (G) above were stimulated with 10 ng/ml hTNF in the presence of zVAD (50 μM) for 24 h followed by Annexin V staining. TNF-induced necroptosis in *Sharpin*$^{-/-}$ cells was reduced by the knockdown of CYLD.

Cell death analyses were conducted with at least three independent experiments. One-way ANOVA analysis was performed on the data set of (A) and (H). Student's t-test was carried out in (B) and (C). Error bars represent mean±SD. *$P<0.05$, **$P<0.01$, ***$P<0.001$, N.S. non-significant.

Supplementary Figure S2

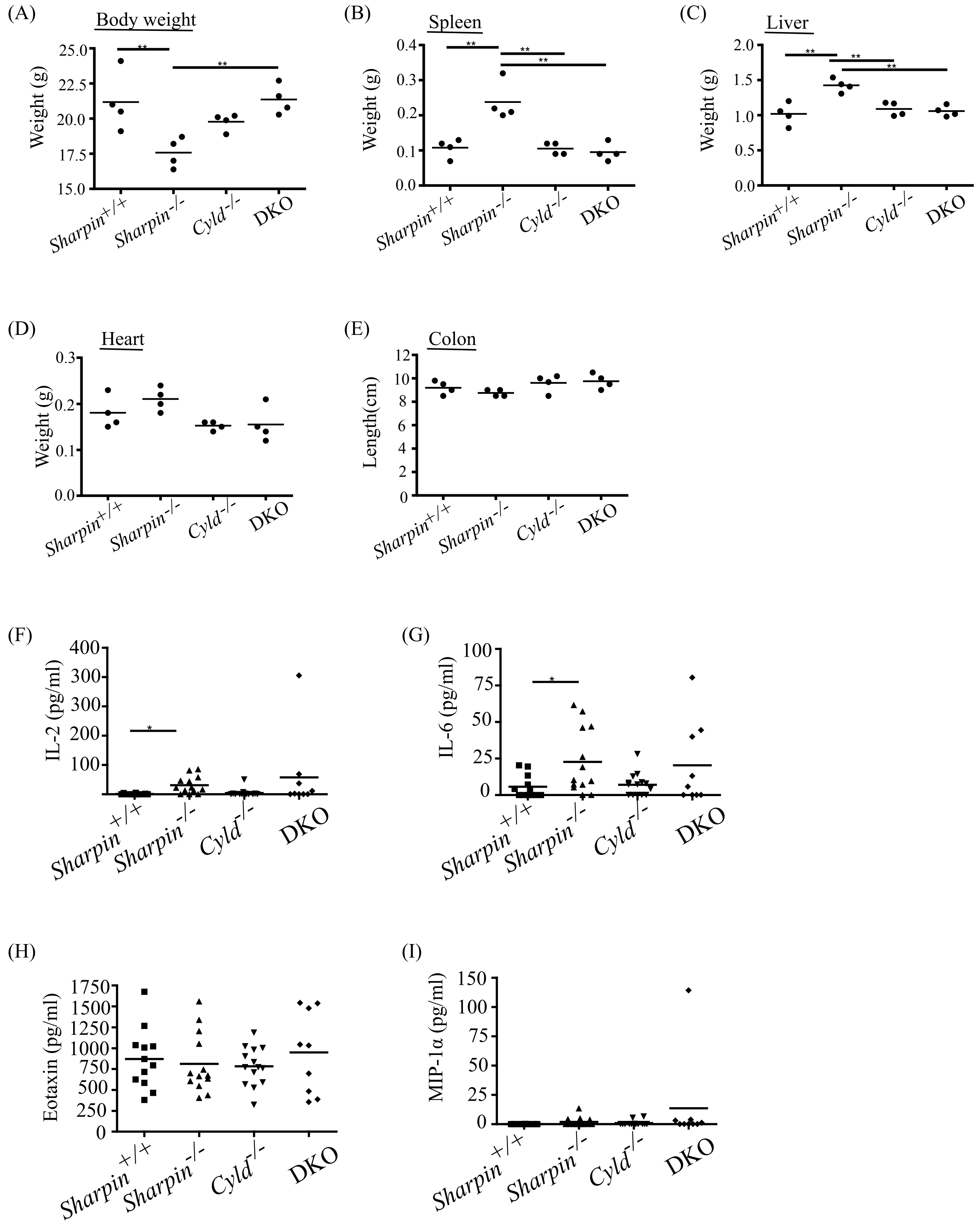

**Supplementary Figure S2. Additional phenotype of *Sharpin*$^{-/-}$*Cyld*$^{-/-}$ DKO mice.**

(A-E) 16-week-old *Sharpin*$^{+/+}$, *Sharpin*$^{-/-}$, *Cyld*$^{-/-}$, and *Sharpin*$^{-/-}$*Cyld*$^{-/-}$ (DKO) female mice were analyzed for the weight of their (A) body (B) spleen (C) liver, (D) heart, and (E) the length of colon. Four mice from each genotype were examined and one-way ANOVA analysis was performed. Error bars represent mean±SD. *P<0.05, **P<0.01. Alterations in body, spleen and liver weight in the *Sharpin*$^{-/-}$ mice were reversed in the *Sharpin*$^{-/-}$*Cyld*$^{-/-}$ DKO.

(F-I) Level of serum (F) IL2, (G) IL6, (H) eotaxin and (I) CCL3 (MIP-1α) from 10-16-week-old *Sharpin*$^{+/+}$, *Sharpin*$^{-/-}$, *Cyld*$^{-/-}$, and DKO mice were measured by Luminex® Cytokines and Chemokines Multiplex Assays. Each data point is from an individual mouse (n=9-12 mice per genotype). The horizontal bar represents median. For panel F & G, data points that were undetermined were denoted as "0" for graph plotting and Mann Whitney U test was performed. Student's t-test was performed on dataset for panel H & I. *P<0.05.

Supplementary Figure S3

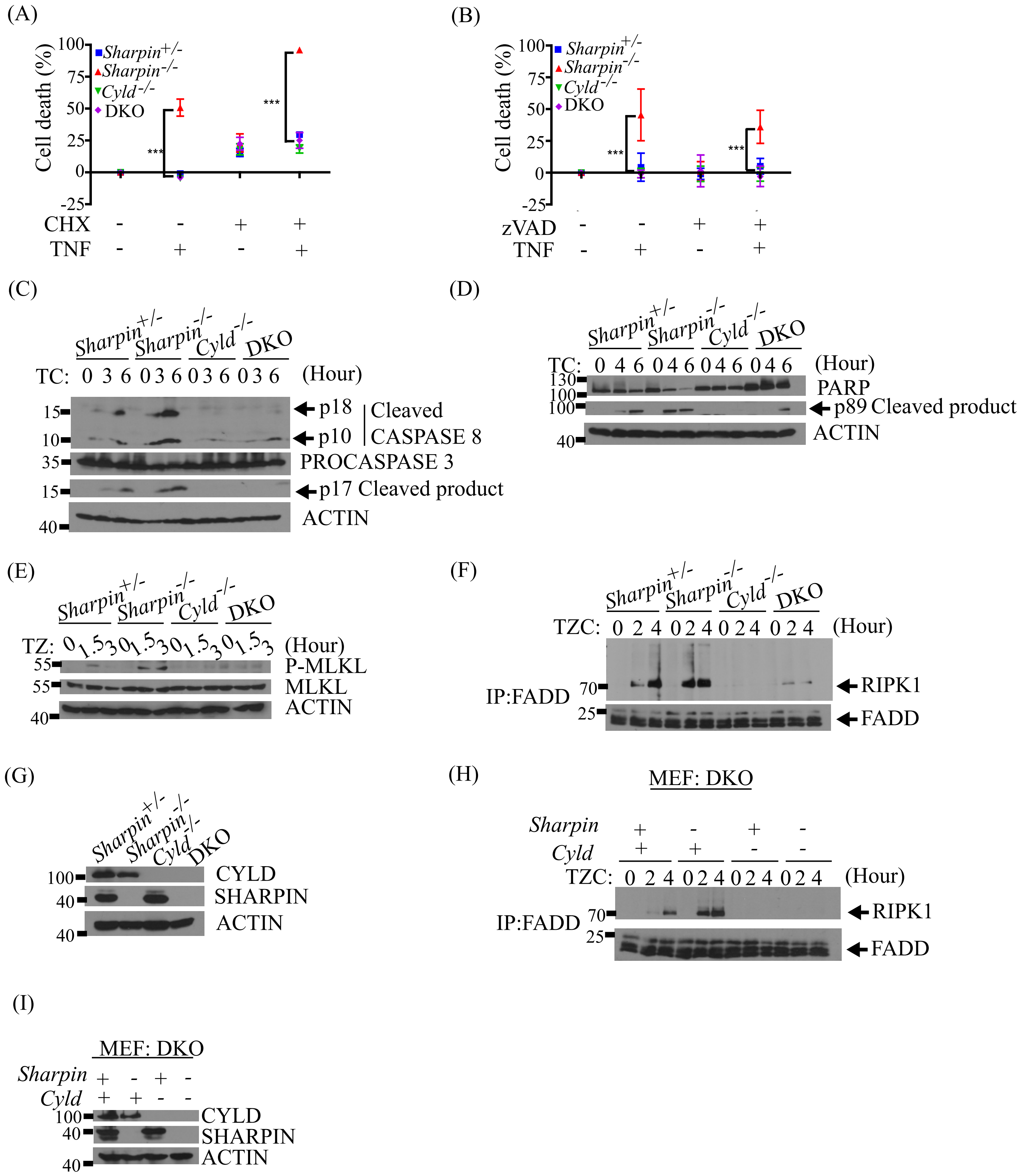

**Supplementary Figure S3.** *Sharpin*$^{-/-}$ MEF are more susceptible to CYLD-mediated cell death.

(A-B) MEF derived from *Sharpin*$^{+/-}$, *Sharpin*$^{-/-}$, *Cyld*$^{-/-}$ and DKO mice were treated for 24h with mTNF (10 ng/ml) in the presence of cycloheximide (CHX, 1 μg/ml) to induce apoptosis or zVAD (20 μM) to induce necroptosis. Cell death was quantified using Cell TiterGlo. At least three independent experiments were conducted and analyzed by one-way ANOVA. Error bars represent mean±SD. *P<0.05, **P<0.01, ***P<0.001. Both forms of death were enhanced in the *Sharpin*$^{-/-}$ cells but were reversed in the DKO cells.

(C-D) *Sharpin*$^{+/-}$, *Cyld*$^{-/-}$, *Sharpin*$^{-/-}$ and DKO MEF were stimulated with mTNF (T, 100 ng/ml) in the presence of cycloheximide (C, 1 μg/ml) for 3 and 6 h. Apoptosis was examined by blotting for cleaved CASP8 and cleaved CASP3 (C) or cleaved PARP-1 (D). *Sharpin*$^{-/-}$ cells exhibited more CASP8, CASP3 and PARP-1 cleavage that were reversed in the DKO cells. Data is representative of three independent experiments.

(E) Necroptosis was analyzed by blotting for phospho-MLKL in *Sharpin*$^{+/-}$, *Sharpin*$^{-/-}$, *Cyld*$^{-/-}$ and DKO MEF after stimulation with mTNF (T, 100 ng/ml) in the presence of zVAD (Z, 20 μM) for 1.5 and 3 h. The enhanced phospho-MLKL observed in the *Sharpin*$^{-/-}$ cells was reversed in the DKO cells. Data is representative of three independent experiments.

(F) *Sharpin*$^{+/-}$, *Sharpin*$^{-/-}$, *Cyld*$^{-/-}$, and DKO MEF were stimulated with mTNF (T, 100 ng/ml) in the presence of cycloheximide (C, 1 μg/ml) and zVAD (Z, 20 μM) for 2 and 4 h. Lysates were subjected to FADD immunoprecipitation to isolate the death-inducing signaling complex (DISC) followed by blotting for RIPK1. There was more translocation of RIPK1 to the DISC in SHARPIN-deficient cells indicating that SHARPIN normally inhibits this process. Furthermore, the RIPK1-FADD DISC formation was dependent on CYLD as it was abrogated in the DKO cells. Data is representative of three independent experiments.

(G) CYLD and SHARPIN expression in the MEF of the four genotypes used in A-F were confirmed by blotting. Data is representative of three independent experiments.

(H) To confirm that the observations in (F) were due to SHARPIN and CYLD, we reconstituted the DKO MEF with SHARPIN alone, CYLD alone, or both SHARPIN and CYLD. An experiment similar to that in (F) was performed with 2 and 4 h stimulation. The results obtained were similar to that in (F). Data is representative of three independent experiments.

(I) Blotting of the reconstituted DKO MEF used in (H) with the indicated antibodies confirmed their complementation with their respective genes. Data is representative of two independent experiments.

Supplementary Figure S4

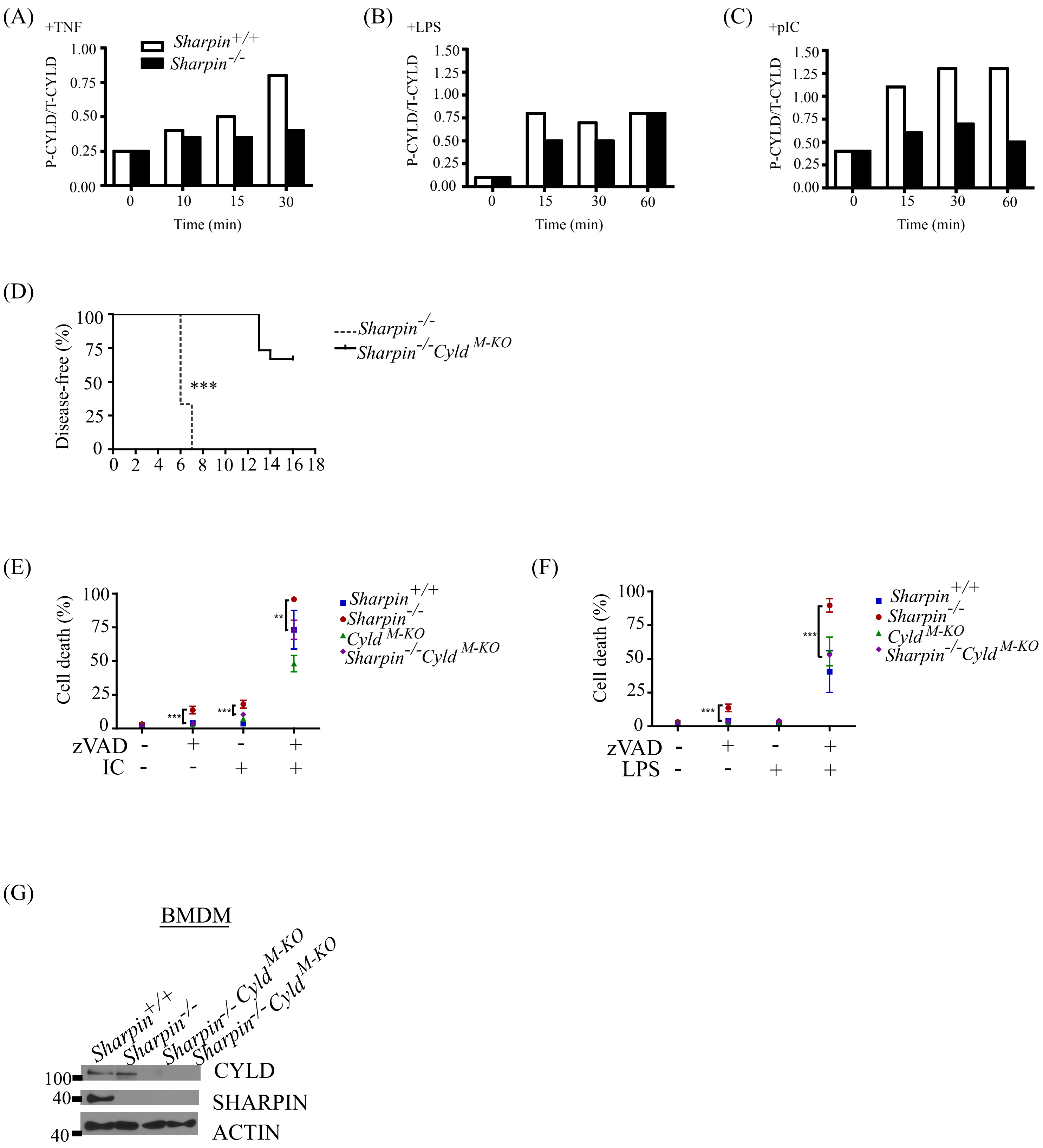

**Supplementary Figure S4. Phosphorylation and function of CYLD in *Sharpin$^{-/-}$* BMDM.**

(A-C) Semi-quantitative analysis of the western blots shown in Figure 5 A-C was conducted using Image J. Values on the Y-axis is the ratio of the signal of the pS418-CYLD band relative to the signal of the total CYLD band.

(D) Kaplan-Meier plot of *Sharpin$^{-/-}$ and Sharpin$^{-/-}$Cyld$^{M-KO}$* cohort exhibiting first signs of dermatitis as determined by appearance of alopecia or excoriations, or 1 small punctate crust i.e., character of lesion =1 as described in Material and Methods (n=10-15 in each group). Student-t test analysis was performed. ***P<0.001.

(E & F) *Sharpin$^{+/+}$*, *Sharpin$^{-/-}$*, *Cyld$^{-/-}$* and *Sharpin$^{-/-}$Cyld$^{M-KO}$* BMDM were treated with 10 μg/ml poly(I:C) (E) or 1 ng/ml LPS (F) for 24 h in the presence or absence of zVAD (20 μM) to induce necroptosis. Cell death was analyzed by propidium iodide staining and flow cytometry. Data from three independent experiments are shown. Error bars represent mean±SD and one-way ANOVA analysis was performed. **P<0.01, ***P<0.001 *Sharpin$^{-/-}$* versus *Sharpin$^{-/-}$Cyld$^{M-KO}$* BMDM. P value of *Sharpin$^{-/-}$Cyld$^{M-KO}$* versus *Sharpin$^{+/+}$* BMDM is not significant.

(G) CYLD and SHARPIN expression in BMDM from *Sharpin$^{+/+}$, Sharpin$^{-/-}$ and Sharpin$^{-/-}$Cyld$^{M-KO}$* were analyzed by blotting to confirm knockout of CYLD in macrophages. BMDM cultures from two individual *Sharpin$^{-/-}$Cyld$^{M-KO}$* mice are shown.

Supplementary Figure S5

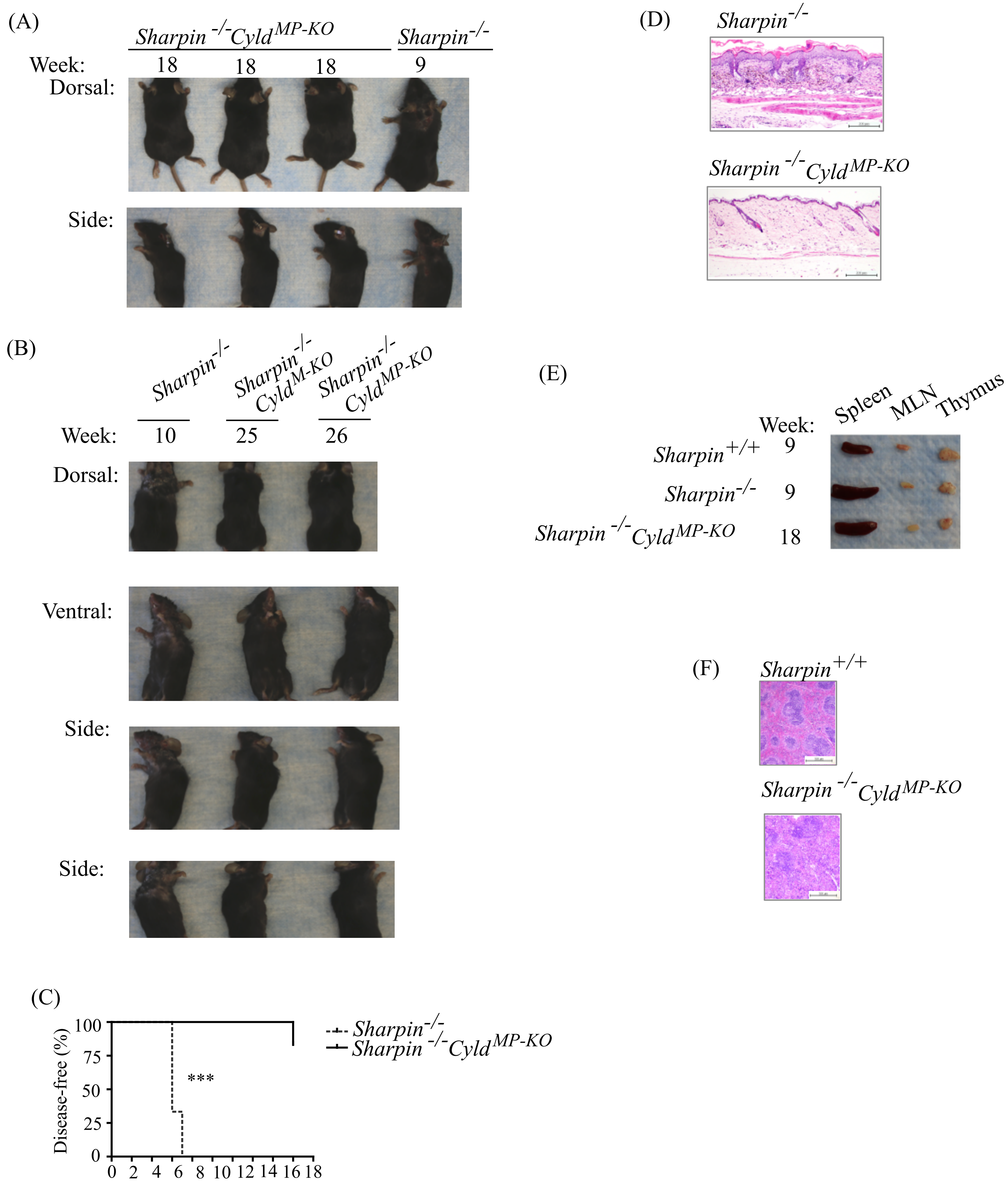

**Supplementary Figure S5. CYLD-mediated death of mononuclear phagocytes is necessary to cause skin inflammation in *Sharpin*$^{-/-}$ mice.**

(A) Photograph of 18-week-old *Sharpin*$^{-/-}$*Cyld*$^{MP\text{-}KO}$ mice in comparison to a 9-week-old *Sharpin*$^{-/-}$ mouse.

(B) Photograph of 25-week-old *Sharpin*$^{-/-}$*Cyld*$^{M\text{-}KO}$ mice, and 26-week-old *Sharpin*$^{-/-}$*Cyld*$^{MP\text{-}KO}$ mice in comparison to a 10-week-old *Sharpin*$^{-/-}$ mouse.

(C) Kaplan-Meier plot of *Sharpin*$^{-/-}$ *and Sharpin*$^{-/-}$*Cyld*$^{MP\text{-}KO}$ cohort exhibiting first signs of dermatitis as described in Supplemental Figure 4D (n=10-15 in each group). Student-t test analysis was performed. ***P<0.001.

(D) H&E staining of skin sections from 16-week-old *Sharpin*$^{-/-}$ and 18-week-old *Sharpin*$^{-/-}$*Cyld*$^{MP\text{-}KO}$ mice (100X).

(E) Gross analysis of spleens, mesenteric lymph nodes and thymi from 18-week-old *Sharpin*$^{+/+}$, *Sharpin*$^{-/-}$ *and Sharpin*$^{-/-}$*Cyld*$^{MP\text{-}KO}$ mice.

(F) H&E staining of spleen sections from 18-week-old *Sharpin*$^{+/+}$ *and Sharpin*$^{-/-}$*Cyld*$^{MP\text{-}KO}$ mice.
